# CAMSAP2 organizes a γ-tubulin-independent microtubule nucleation centre

**DOI:** 10.1101/2021.03.01.433304

**Authors:** Tsuyoshi Imasaki, Satoshi Kikkawa, Shinsuke Niwa, Yumiko Saijo-Hamano, Hideki Shigematsu, Kazuhiro Aoyama, Kaoru Mitsuoka, Mari Aoki, Ayako Sakamoto, Yuri Tomabechi, Naoki Sakai, Mikako Shirouzu, Shinya Taguchi, Yosuke Yamagishi, Tomiyoshi Setsu, Yoshiaki Sakihama, Takahiro Shimizu, Eriko Nitta, Masatoshi Takeichi, Ryo Nitta

## Abstract

Microtubules are dynamic polymers consisting of αβ-tubulin heterodimers. The initial polymerization process, called microtubule nucleation, occurs spontaneously via αβ-tubulin. Since a large energy barrier prevents microtubule nucleation in cells, the γ-tubulin ring complex is recruited to the centrosome to overcome the nucleation barrier. However, detachment of a considerable number of microtubules from the centrosome is known to contribute to fundamental processes in cells. Here, we present evidence that minus-end-binding calmodulin-regulated spectrin-associated protein 2 (CAMSAP2) serves as a strong nucleator for microtubule formation from soluble αβ-tubulin independent of γ-tubulin. CAMSAP2 significantly reduces the nucleation barrier close to the critical concentration for microtubule polymerization by stabilizing the longitudinal contacts among αβ-tubulins. CAMSAP2 clusters together with αβ-tubulin to generate nucleation intermediates, from which numerous microtubules radiate, forming aster-like structures. Our findings suggest that CAMSAP2 supports microtubule growth by organizing a nucleation centre as well as by stabilizing microtubule nucleation intermediates.

## Introduction

Microtubules are dynamic tubular polymers that contribute to fundamental processes in cells, such as cell shape determination, chromosome segregation, cilium and flagellum formation, and molecular motor trafficking (Desai and Mitchison, 1997). Microtubules are composed of αβ-tubulin heterodimers. α-Tubulin and β-tubulin align in a head-to-tail manner to form protofilaments that associate laterally to grow into microtubules (Alushin et al., 2014; Nogales et al., 1999). The exposed β-tubulin end is called the plus-end, whereas the other side, the exposed α-tubulin end, is called the minus-end. The microtubule plus-end is more dynamic than the minus-end; it dynamically alternates between growth and shrinkage. The dynamic events are regulated by many microtubule plus-end-tracking proteins (+TIPs) (Akhmanova and Steinmetz, 2015, 2008; Howard and Hyman, 2003). The minus-end is less dynamic and is responsible for determining the geometry of microtubule networks by stably anchoring at microtubule nucleation sites or the microtubule organizing centre (Akhmanova and Steinmetz, 2019; Dammermann et al., 2003).

Despite the identification of many +TIPs, only a few minus-end-binding proteins (- TIPs) have been reported, including γ-Tubulin and calmodulin-regulated spectrin-associated proteins (CAMSAPs). γ-Tubulin is located at centrosomes to nucleate microtubules and stabilize the microtubule minus-ends (Akhmanova and Steinmetz, 2015). γ-Tubulin forms a ring complex (γ-TuRC), which serves as a nucleation template for centrosomal microtubule formation by aligning αβ-tubulins to form the tubular structure of the microtubule in a process called templated nucleation (Moritz et al., 1995; Roostalu and Surrey, 2017; Wieczorek et al., 2020; Zheng et al., 1995). CAMSAPs, the protein family composed of CAMSAP1-3 in vertebrates, Patronin in *Drosophila melanogaster*, and PaTRoNin (microtubule-binding protein) homolog (PTRN-1) in *Caenorhabditis elegans*, have been reported to be involved in non-centrosomal microtubule formation (Goodwin and Vale, 2010; Marcette et al., 2014; Meng et al., 2008; Richardson et al., 2014; Wu et al., 2016). Although several *in vitro* reconstitution experiments have shown differences among CAMSAPs, CAMSAPs generally bind to the growing microtubule minus-ends and strongly suppress the dynamicity of non-centrosomal microtubules in different cell types, including neurons and epithelial cells (Chuang et al., 2014; Hannak et al., 2002; Jiang et al., 2014; Martin et al., 2018; Nashchekin et al., 2016; Noordstra et al., 2016; Pongrakhananon et al., 2018; Sampaio et al., 2001; Tanaka et al., 2012; Toya et al., 2016; Wu et al., 2016; Yau et al., 2014). The minus-end side of the microtubule lattice, but not the very end, is recognized by the CKK domain, although some additional effects were detected by other regions of CAMSAPs (Atherton et al., 2019, 2017). CAMSAPs have been reported to act in parallel with γ-TuRC-dependent microtubule nucleation, and the microtubule-severing protein katanin regulates the functions of CAMSAPs (Jiang et al., 2018; Wang et al., 2015; Wu et al., 2016). CAMSAPs are thought to be rapidly recruited to and to decorate pre-existing nascent microtubule minus-ends to sustain non-centrosomal microtubules. On the other hand, some reports have demonstrated the possibility that CAMSAP-containing foci are involved in promoting microtubule nucleation independently of γ-TuRC (Atherton et al., 2019; Jiang et al., 2018; Nashchekin et al., 2016) Thus, the mechanism by which CAMSAP2 proteins contribute to non-centrosomal microtubule organization has remained controversial.

Recent studies suggest that microtubules in cells often nucleate independently of the γ-TuRC template (Hannak et al., 2002; O’Toole et al., 2012; Sampaio et al., 2001; Yau et al., 2014). Therefore, we examined the roles of CAMSAPs in microtubule nucleation and microtubule network formation *in vitro*. In this study, we discovered the fundamental role of CAMSAP2 as a strong nucleator of microtubule formation from soluble αβ-tubulins. Our structural studies clearly demonstrated that CAMSAP2 co-condenses with tubulins to stimulate spontaneous nucleation of microtubules without the γ-TuRC template. Many microtubule nucleation intermediates, tubulin rings, tubulin sheets, and their mixtures were observed inside the CAMSAP2-tubulin condensate during the early stages of microtubule polymerization. The CAMSAP2-tubulin condensate grows into the nucleation centre and mediates aster-like microtubule network formation *in vitro*, demonstrating the importance of CAMSAP2 in non-centrosomal, γ-tubulin-independent microtubule nucleation processes.

## Results

### Profiles of CAMSAP2 constructs used in *in vitro* biochemical, biophysical, and structural assays

Before performing functional analyses of CAMSAP2, we examined the biochemical properties of the recombinant CAMSAP2 constructs used in this study by SDS-PAGE and size exclusion chromatography (Figure 1A). As shown in Figure 1B and C, full length CAMSAP2 with and without GFP-tag constructs were highly purified and showed a sharp symmetrical single peak in size exclusion chromatography, suggesting their homogeneous size distribution in solution. GFP-tagged full-length was eluted slightly faster than the non-tagged construct due to the size of GFP-tag, indicating both constructs should be in the same oligomeric state in solution.

**Figure 1.**
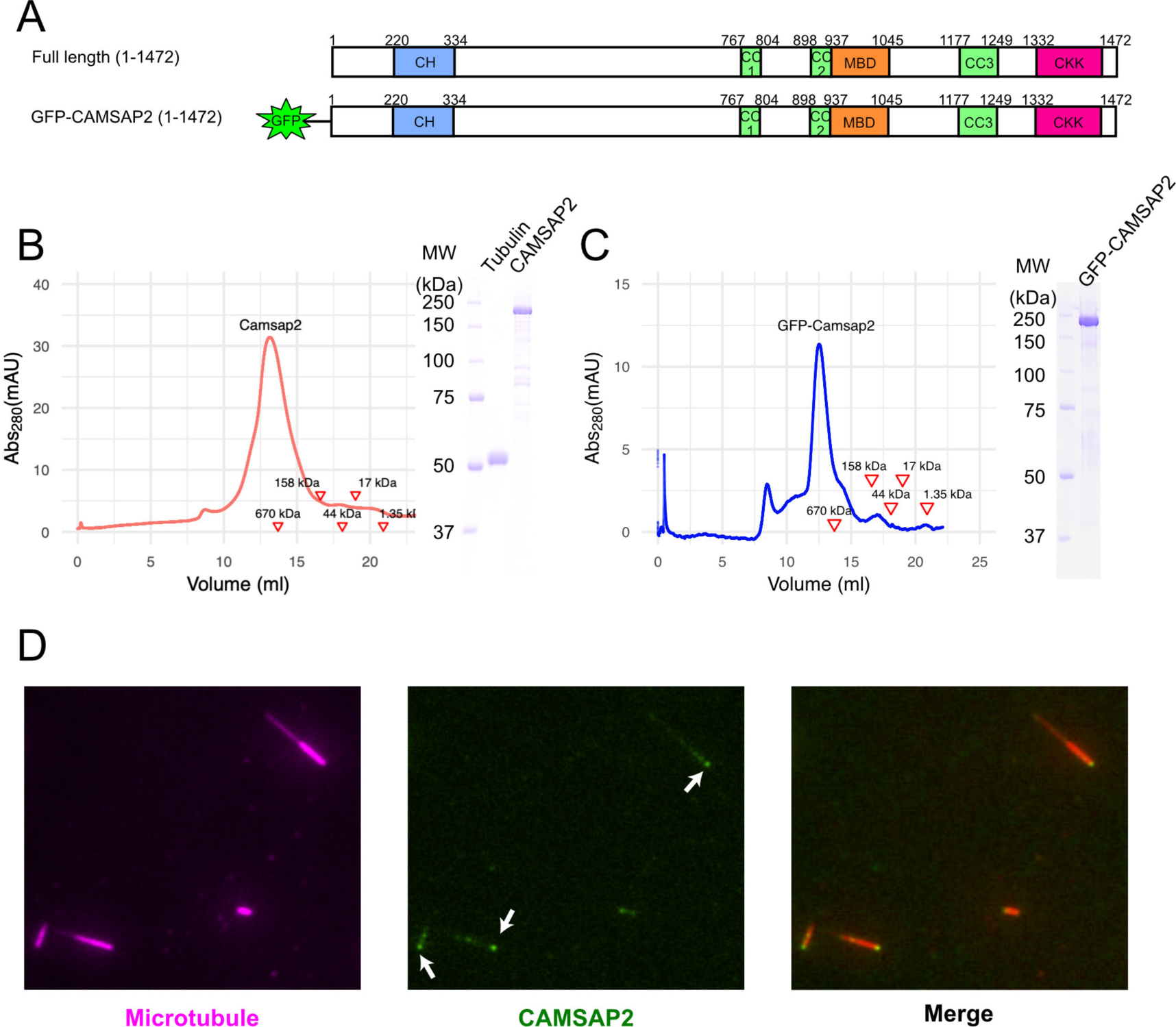
Functional study of recombinant CAMSAP2. (A) Schematic diagram of the full lengthCAMSAP2 constructs used in this study. CH: calponin-homology domain; MBD: microtubule-binding domain; CC: coiled-coil domain; CKK: C-terminal domain common to CAMSAP1 and two other mammalian proteins, KIAA1078 and KIAA1543. (B)(C) Size exclusion chromatography and SDS-PAGE of the peak fraction of (B) full-length CAMSAP2 and (C) GFP-CAMSAP2. (D) TIRF images of polarity-marked microtubules (magenta) decorated with purified GFP-CAMSAP2 (green). The minus-end segment of the microtubule is brighter than the plus-end segment.

We then analysed the microtubule binding pattern of recombinant full-length GFP-CAMSAP2 by TIRF microscopy. The results showed that CAMSAP2 bound to the microtubule lattice preferentially via the minus-ends, consistent with previous results (Atherton et al., 2017; Jiang et al., 2014) (Figure 1D). These results collectively indicate that the CAMSAP2 proteins used in this study retained their original ability to bind to microtubule minus-ends.

### Microtubule nucleation with CAMSAP2 occurs at approximately the critical concentration for microtubule polymerization

Microtubule nucleation from αβ-tubulin heterodimers (hereafter called tubulin in the Results section) can occur spontaneously *in vitro* (Kuchnir Fygenson et al., 1995; Voter and Erickson, 1984). *De novo* microtubule nucleation *in vitro* requires a critical tubulin concentration greater than 20 µM. This concentration is considerably higher than that required for elongation from existing microtubules, which is typically approximately 1-2 µM and is called the critical concentration for microtubule polymerization (Cc_MT polymerization_) (Wieczorek et al., 2015).

To test the nucleation ability of the purified pig tubulins used in this study, we performed a spin-down spontaneous nucleation assay through microtubule co-pelleting experiments *in vitro* (Wieczorek et al., 2015). Microtubule nucleation was induced by 60 min of incubation at 37 °C with high-speed centrifugation. To exclude non-specific aggregation during the microtubule pelleting assay, the microtubule pellets were resuspended and depolymerized on ice for 30 min. After incubation on ice, the samples were centrifuged again, and the supernatant was applied for SDS-PAGE analysis (Figure 2A). *De novo* nucleation of microtubules was detected just above the tubulin concentration of 20 µM, which is consistent with a previous report (Wieczorek et al., 2015) (Figure 2A and C). The ratio of polymerized microtubules to unpolymerized tubulin increased proportionately with the tubulin concentrations.

**Figure 2.**
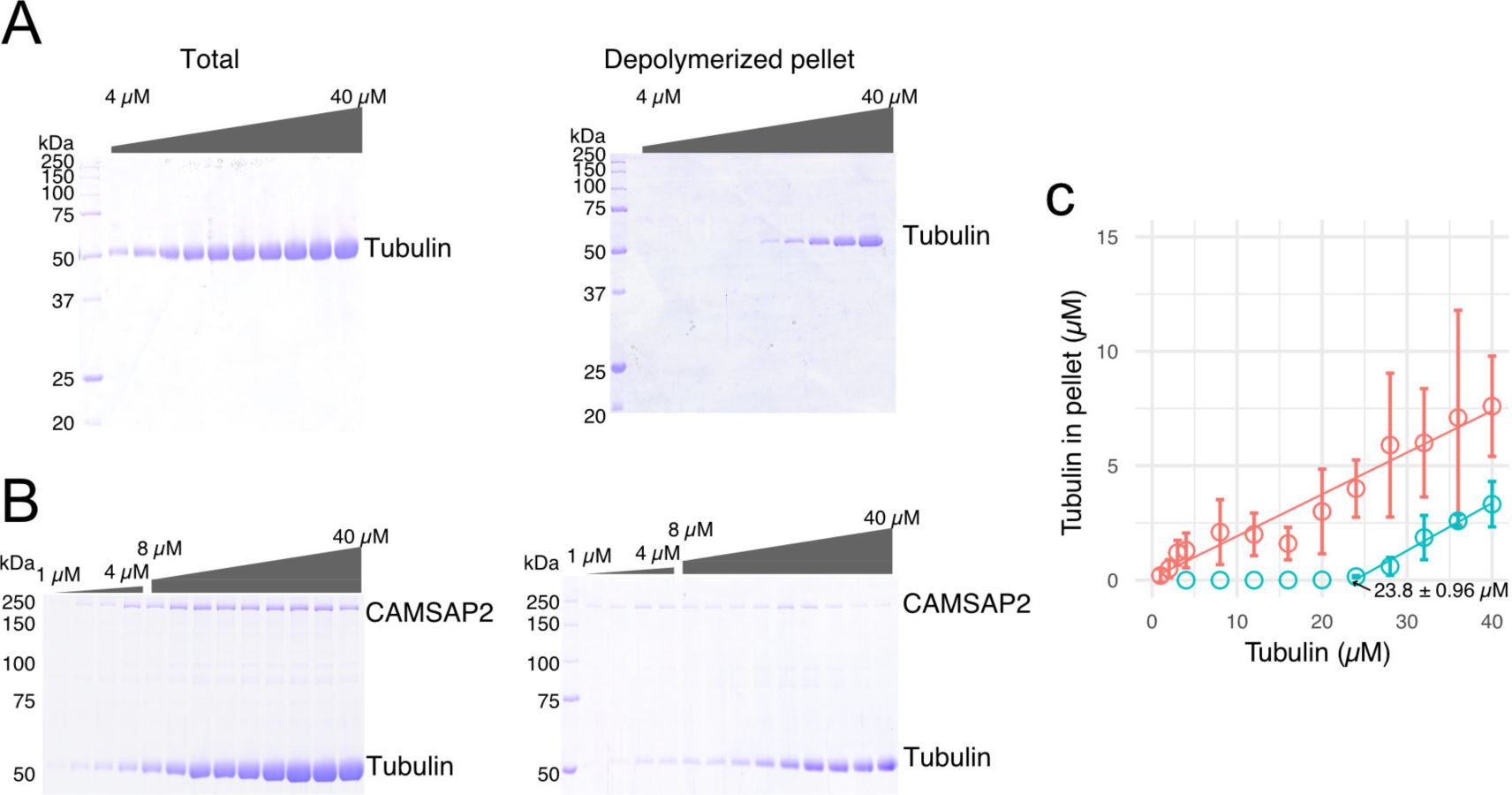
CAMSAP2 stimulates microtubule nucleation. (A) SDS-PAGE gels from a spin-down spontaneous nucleation assay of tubulin showing the total tubulin after 60 min of polymerization at 37 °C (left gel). Polymerized tubulin was pelleted by centrifugation and then depolymerized on ice and centrifuged to remove debris, and the supernatant was subjected to SDS-PAGE (right gel). (B) SDS-PAGE gels from a spin-down spontaneous nucleation assay of tubulin with 1 µM CAMSAP2 showing the total tubulin-CAMSAP2 (left gel) and the polymerized/depolymerized tubulin (right gel). (C) Plots of the depolymerized tubulin concentrations determined by pelleting assay against the total tubulin concentrations determined by reaction on SDS-PAGE gel from three independent assays. The depolymerized tubulin from the tubulin assay is turquoise green, and that combined with CAMSAP2 is orange. The concentrations of tubulin greater than 0.1 µM are fitted with a trend line that has an x-intercept of 23.8 ± 0.96 µM.

We then evaluated the effect of CAMSAP2 on microtubule nucleation by co-pelleting assay. Both tubulin and CAMSAP2 were pelleted from 1.0-2.0 µM, suggesting that tubulins in the presence of CAMSAP2 could polymerize into microtubules at 1.0-2.0 µM, which is very close to the Cc_MT polymerization_ (Figure 2B and C). This result suggests that CAMSAP2 drastically reduced the nucleation barrier close to the theoretical limit of microtubule polymerization. In this assay, both tubulin and CAMSAP2 in the pellet were recovered to the supernatant after depolymerization on ice, indicating that CAMSAP2 co-precipitates with microtubules. Similar to the behaviour of microtubule polymerization and depolymerization, CAMSAP2 forms a reversible protein complex cluster with microtubules. Based on these results, we concluded that CAMSAP2 significantly enhances the nucleation of tubulin by forming a protein complex cluster with tubulins or microtubules.

### CAMSAP2 and tubulin co-condense to form aster-like structure

To investigate how CAMSAP2 affects the nucleation and polymerization of microtubules, we co-polymerized 10 µM of tubulin with CAMSAP2 and observed them by fluorescence microscopy. Tetramethylrhodamine (TMR, 10 μM)-labelled tubulin was co-incubated with 10 to 1,000 nM full-length GFP-CAMSAP2 at 37 °C for 10 min (Figure 3A). Surprisingly, CAMSAP2 co-condensed with tubulin and microtubule bundles extended radially outward from the condensates (Figure 3B). Their shapes were reminiscent of the centrosomal aster; thus, we named them as “Cam2-asters”. Cam2-asters appeared from 250 nM CAMSAP2, and the number of asters increased as the CAMSAP2 concentration increased from 250 to 1,000 nM CAMSAP2 (Figure 3B and C). CAMSAP2 localized mainly at the central regions of Cam2-asters.

**Figure 3.**
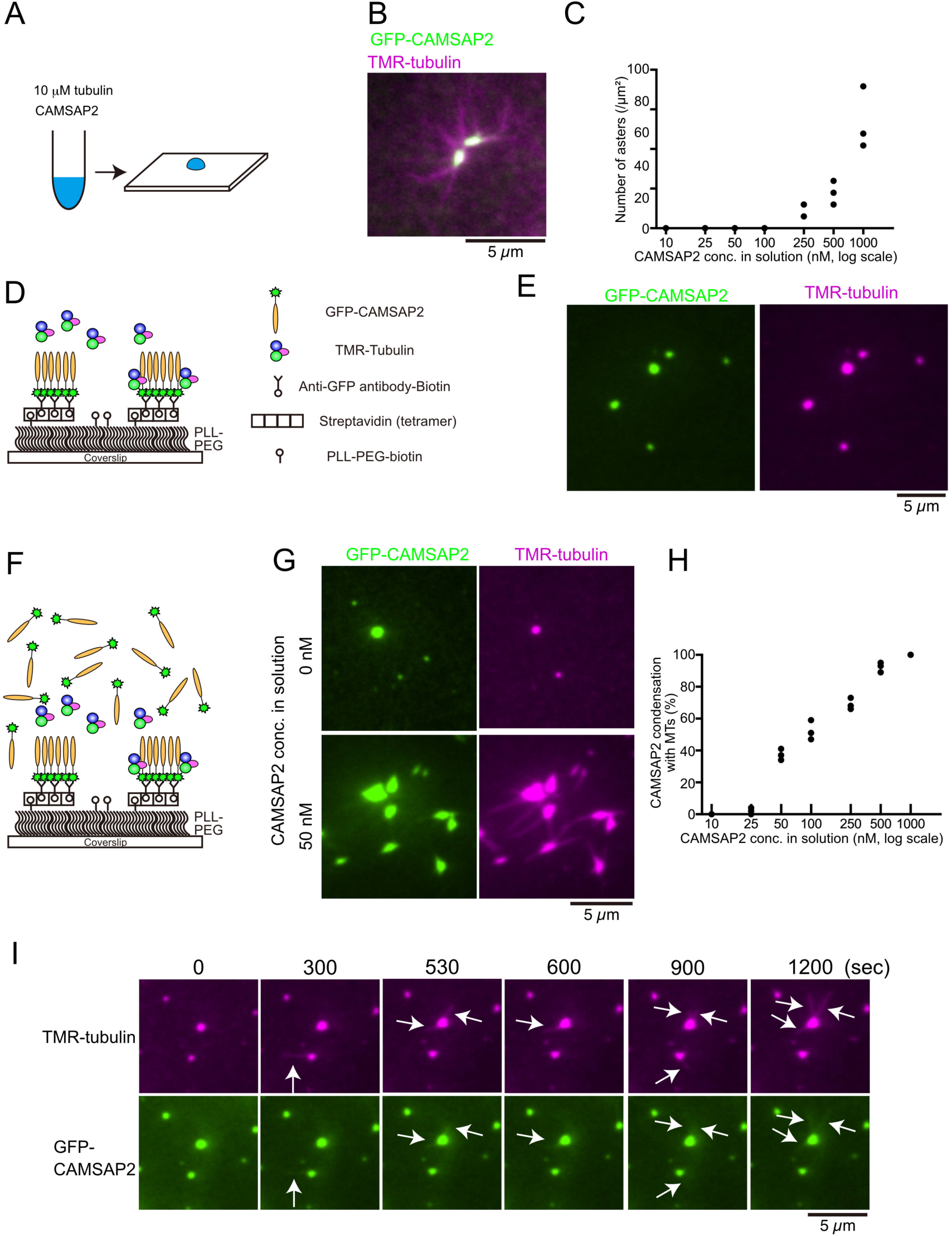
Tubulin is incorporated into CAMSAP2 clusters to form aster-like structure. (A) Procedure used to obtain the data in panels (B) and (C). GFP-CAMSAP2, 10 μM tubulin and 0.5 μM TMR-tubulin were mixed in BRB80 supplemented with 100 mM KCl and incubated for 10 min at 37 °C. The solution was directly transferred onto a coverslip and observed by fluorescence microscopy. (B) Representative image of asters. GFP-CAMSAP2 (0.5 μM), tubulin (10 μM) and TMR-tubulin (0.5 μM) were co-incubated. (C) Quantification of the numbers of asters in solutions containing 10 μM tubulin, 0.5 μM TMR-tubulin and 10, 25, 50, 100, 250, 500, and 1000 nM CAMSAP2. The results of three independent assays are shown with dots. (D) Schematic showing reconstitution of CAMSAP2-containing foci. (E) Soluble tubulins were incorporated into CAMSAP2 condensates within 1 min. (F) Schematic showing CAMSAP2-containing foci with soluble tubulin and CAMSAP2. (G) CAMSAP2 in solution induced aster formation from CAMSAP2 condensates in a dose-dependent manner. Representative images for 0 and 50 nM CAMSAP2 are shown. (H) Quantification of microtubule formation from CAMSAP2 condensates. The percentages of CAMSAP2 condensates with microtubules among total CAMSAP2 condensates are shown. Each dot shows the results of three independent experiments. (I) Time-lapse images of aster formation. Tubulin (10 μM), TMR-tubulin (0.5 μM) and CAMSAP2 (50 nM) were incubated with CAMSAP2 condensates fixed on coverslips. Dynamic microtubules from CAMSAP2 condensates were observed (arrows). The scale bars indicate 5 µm. See *Videos 1-2*.

**Video 1.** Time-lapse movie of aster formation. Tubulin (10 μM), TMR-tubulin (0.5 μM) and CAMSAP2 (50 nM) were incubated with CAMSAP2 condensates fixed on coverslips. The size of the field is 81.9 μm x 81.9 μm. The frame rate is 0.2 frames/sec.

**Video 2.** Magnified movie of aster formation from 1. Dynamic microtubules from CAMSAP2 condensates were clearly observed. The size of the field is 8.2 μm x 8.2 μm. The frame rate is 0.2 frames/sec.

The CAMSAP2 concentration required for condensates formation *in vitro* was slightly higher than the physiological concentration of CAMSAP2 (Liebermeister and Klipp, 2006; Wühr et al., 2014). Previous reports have indicated that CAMSAPs are not only cytosolic but also concentrated at sub-cellular structures such as adherens junctions and the apical cortices of epithelial cells (Toya et al., 2016). These CAMSAP-containing foci have been suggested to be involved in promoting microtubule nucleation. To test this hypothesis, we reconstituted CAMSAP-containing foci *in vitro* using a streptavidin-biotin system (Figure 3D). In this system, theoretically, a maximum of three CAMSAP2 dimers were anchored to one biotin molecule that was attached to a coverslip. TMR-labelled tubulin was added to the CAMSAP2-containing foci and visualized by TIRF microscopy. Tubulin was efficiently incorporated into the CAMSAP2 foci within 1 min, although polymerization of microtubules was rarely observed (Figure 3E). We hypothesized that microtubule polymerization from CAMSAP2 foci may also require free CAMSAP2. To test this hypothesis, we supplemented the buffer with 10 to 1,000 nM soluble CAMSAP2 (Figure 3F). As a consequence, microtubule formation was induced from the CAMSAP2 foci. CAMSAP2 at a concentration of 50 nM in solution was sufficient for efficient microtubule polymerization from the foci, and in the presence of 500 nM CAMSAP2, microtubule polymerization occurred at nearly 100% of foci (Figure 3G, H).

Next, we acquired time-lapse images of microtubules using GFP-labelled CAMSAP2 and TMR-labelled tubulin by TIRF microscopy. We found that dynamic polymerization and depolymerization of microtubules occurred at CAMSAP2-containing foci (Figure 3I, Videos 1 and 2). Closer observations also indicated that CAMSAP2 was associated with these dynamic microtubules. These findings suggest that fixed foci of CAMSAP2 act as cores for microtubule polymerization, leading to the formation of Cam2-asters, and that soluble CAMSAP2 is also required for this process. Thus, it seems that CAMSAP2 plays a dual role, functioning as a microtubule-organizing centre and increasing the efficiency of microtubule polymerization.

### Functional domain mapping for CAMSAP2 microtubule-nucleation and Cam2-aster formation

To visualize the further detail of Cam2-aster formation, we co-polymerized tubulin and CAMSAP2 and observed them by negative stain electron microscopy (EM). 10 µM Tubulin was incubated with 1 µM CAMSAP2 at 37 °C for 30 min. Consistent with the results of the fluorescence microscopy assay described above, we observed that spherical shapes of condensates distributed throughout the grid (Figure 4A).

**Figure 4.**
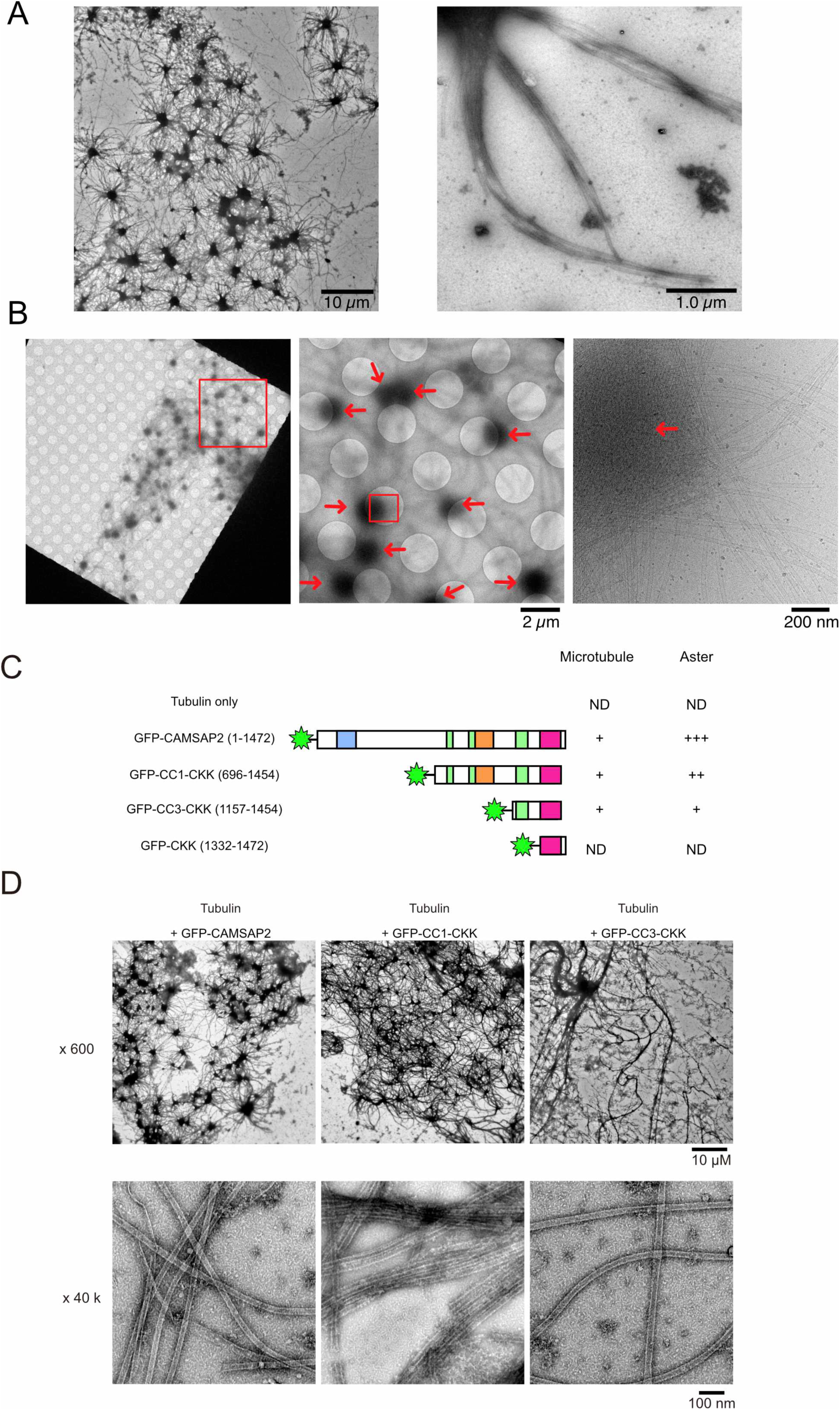
Functional domain mapping of the microtubule nucleation and aster formation activity of CAMSAP2. (A) Negative stain EM micrographs of 10 µM tubulin polymerized with 1 µM of CAMSAP2 after incubation at 37 °C for 10 min. Aster-like microtubule structures were observed. The scale bars indicate 10 µm and 1 µm. (B) Cryo-EM micrographs of 30 µM tubulin polymerized with 3 µM CAMSAP2 after incubation at 37 °C for 10 min captured at different magnifications. Cam2-asters are indicated by the red arrows. (C) Microtubule nucleation and aster formation activities of CAMSAP2 deletion constructs evaluated by the results of 10 µM tubulin with 1 µM CAMSAP2. The number of “+” symbols indicate the strength of the activity (+++: strongest; +: weakest; ND: not detected). Size exclusion chromatography and SDS-PAGE of GFP fused constructs are available in Figure 4—figure supplement 1. (D) Negative stain EM images of polymerization by 10 µM tubulin with 1 µM full-length CAMSAP2 or 1 µM CAMSAP2 mutants during 10 min of incubation at 37 °C. The results for tubulin alone and GFP-CKK are available in Figure 4—figure supplement 2.

**Figure 4—figure supplement 1.**
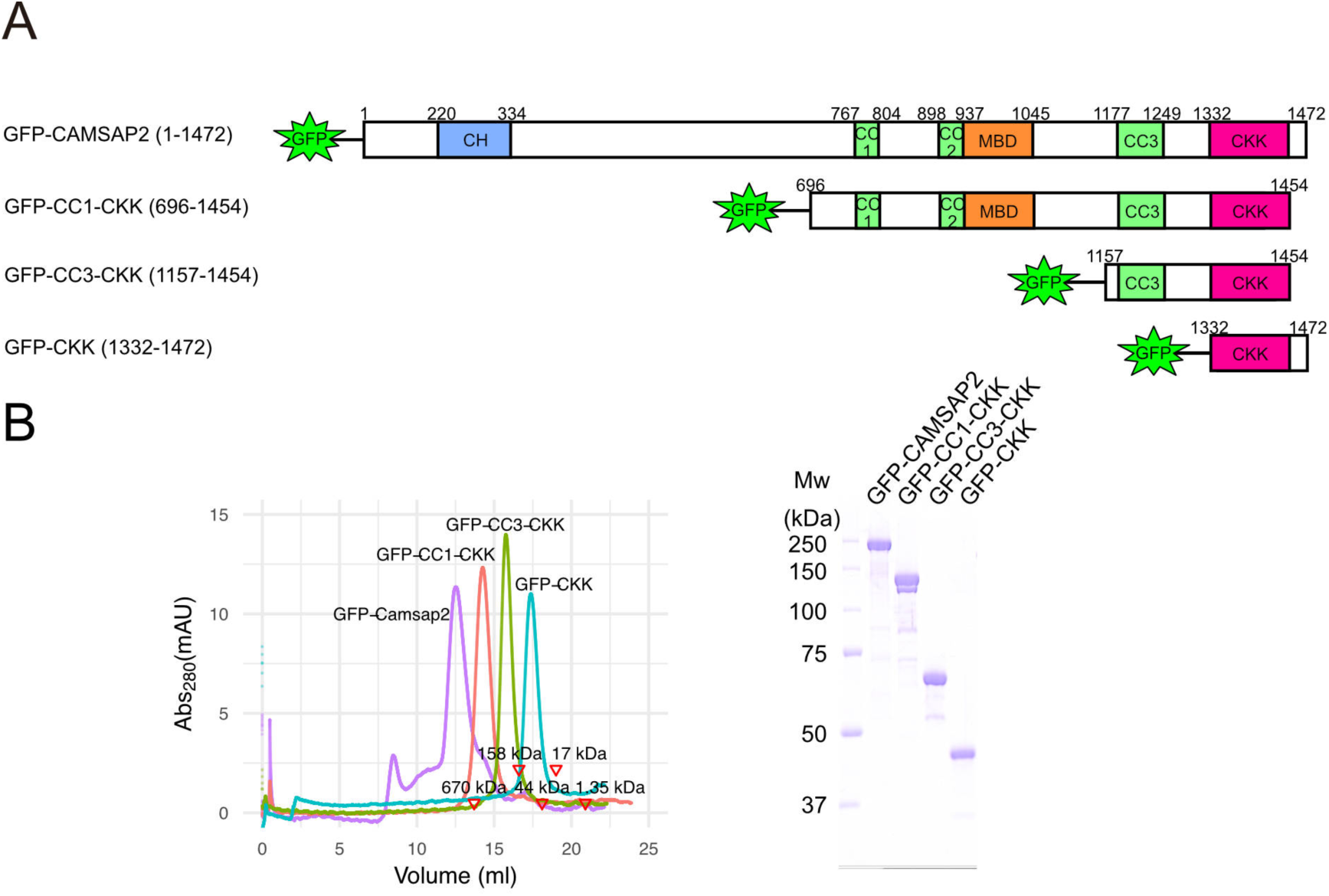
GFP fused CAMSAP2 constructs used in this study. (A) Schematic diagram of the GFP fused CAMSAP2 constructs used in this study. (B) Size exclusion chromatography of GFP-CAMSAP2 (purple), GFP-CC1-CKK (red), GFP-CC3-CKK (green), and GFP-CKK (cyan). The peaks of each sample were subjected to SDS-PAGE.

**Figure 4—figure supplement 2.**
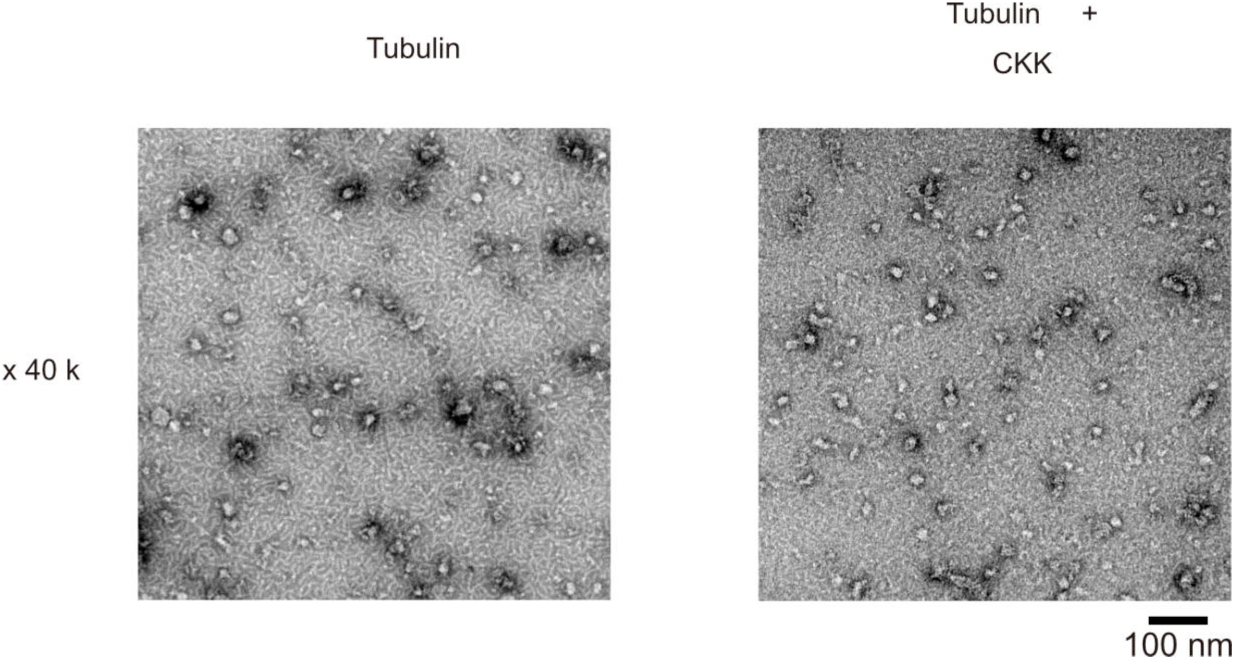
Functional domain mapping of CAMSAP2 analysed by negative stain EM. Ten-micromolar tubulin was incubated alone (left) or with 1 µM GFP-CKK (right) at 37 °C for 10 min and analysed by negative stain EM.

From the condensates, many microtubule bundles extended outward. Microtubule bundles from the neighboring condensates further interacted with each other and formed a dense microtubule meshwork (Figure 4A).

To further eliminate the possible artifacts of Cam2-aster network formation induced by negative staining, we observed the samples by cryo-EM. Consequently, cryo-EM clearly visualized the Cam2-asters (Figure 4B), although the centres of the asters were too thick to be resolved by transmission electron microscopy (TEM) (Figure 4B). From the CAMSAP2 clusters, plenty of microtubule bundles extended outward.

Next, to identify the functional domain of CAMSAP2 for microtubule nucleation and Cam2-aster formation, we prepared a series of N-terminal deletion constructs and tested their functions (Figure 4C and Figure 4—figure supplement 1). Tubulin (10 µM) was mixed with each of the CAMSAP2 mutants (1 µM) on ice, incubated at 37 °C for 10 min, and examined for microtubule nucleation and aster-forming activity (Figure 4C and 4D). No microtubules were observed after 10 min of incubation with tubulin alone (Figure 4—figure supplement 2), consistent with the results of the microtubule pelleting assay (Figure 2). Tubulin in the presence of full-length GFP-CAMSAP2 produced a dense Cam2-aster network as described above (Figure 4D, left), which functions equivalent to the no-GFP CAMSAP2. In the presence of CC1-CKK, the Cam2-aster network was similarly observed, although the condensed region of CAMSAP2-tubulin at the centre of each aster was smaller than that observed with full-length CAMSAP2 (Figure 4D, centre). CC3-CKK could also form Cam2-asters, but condensations were less represented and the meshwork was sparser than observed for the full-length and CC1-CKK constructs (Figure 4D, right). We also examined the copolymerization assay in the presence of CKK domain, which is the major binding site for the microtubule. Consequently, polymerized microtubules and Cam2-asters were rarely observed in the presence of CKK (Figure 4—figure supplement 2). Thus, we concluded only CKK domain could not stimulate tubulin nucleation and Cam2-aster formation. Coiled-coil regions, at least the CC3 domain, are thus required for microtubule polymerization and Cam2-aster formation, which indicates that the CC3 and CKK domains are the minimum requisites for microtubule nucleation and Cam2-aster formation by CAMSAP2.

### 3D architecture of Cam2-aster observed by cryo-electron tomography

To understand the 3D architecture of Cam2-asters, we then performed cryo-electron tomography (cryo-ET) of mature Cam2-asters formed via 30 min of incubation of 10 µM tubulin with 1 µM CAMSAP2 at 37 °C (Figure 5A and B, Video 3). In the tomogram, numerous microtubule bundles were observed to extend outward from the condensates of Cam2-asters. However, we still could not resolve their inner structures because the central regions of the mature Cam2-asters were too thick for the electron beam to penetrate, even with a 300 kV Titan Krios cryo-electron microscope. The distances among the microtubules in the bundle were not always uniform, and no cross-linking density, including that of CAMSAP2, was observed among them, at least at the current resolution (Figure 5A to C). Considering the localization of fluorescence CAMSAP2 at the condensates, CAMSAP2 should have mainly localized at the centre of the aster. Intriguingly, several microtubule ends with ring structures were observed close to the Cam2-aster centre (Figure 5D), as detailed later.

**Figure 5.**
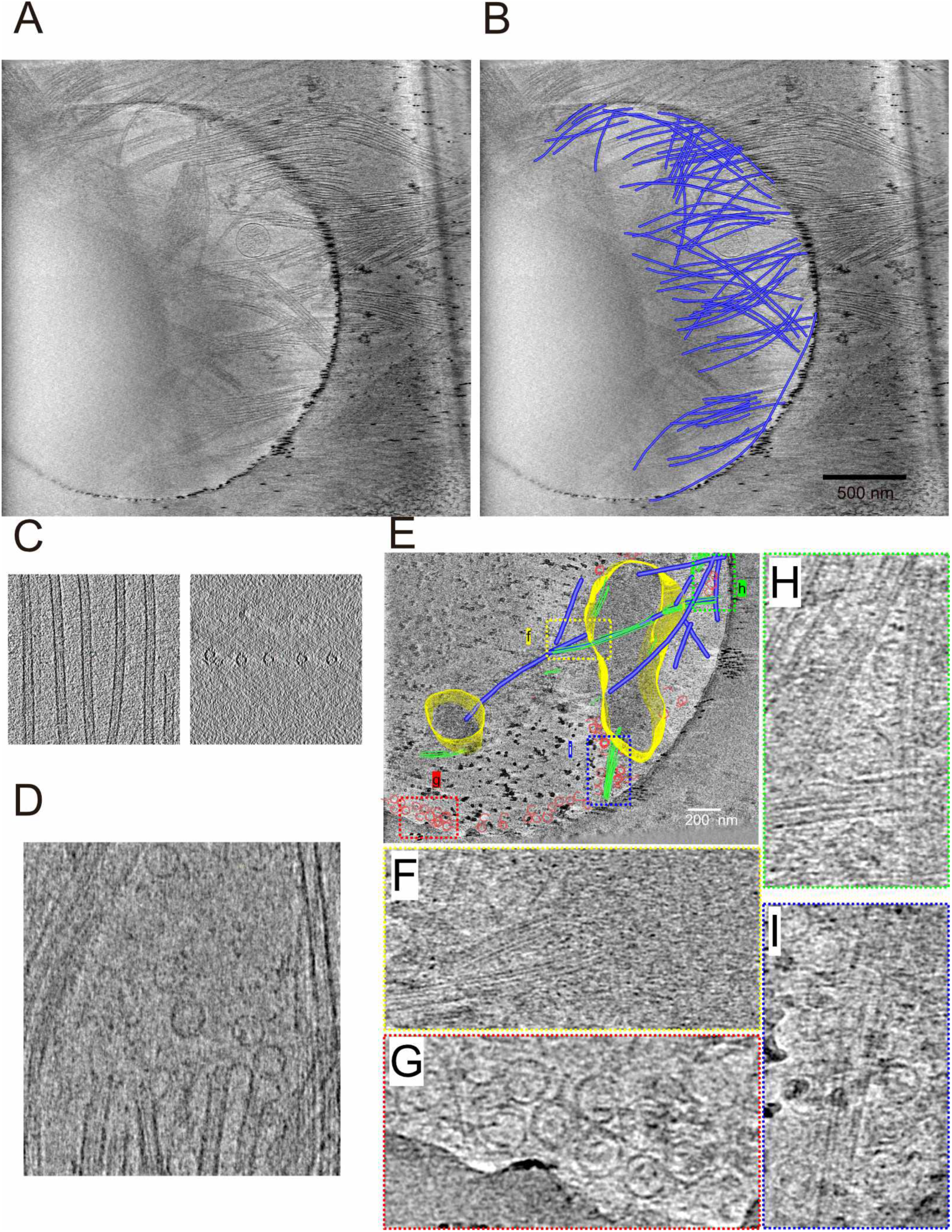
Cryo-ET reconstruction of Cam2-asters. (A) A 2.1 nm thick tomographic slice of a mature Cam2-aster processed by SIRT. See also *Video 3*. (B) Rendered microtubules (blue) from representative 600 nm thick tomographic slices overlaid with a tomographic slice. The centre of the Cam2-aster is located in the left area of the grid hole. (C) Typical views of microtubule bundles elongated from an aster are shown from different angles. (D) Ring structures were observed near the microtubule ends. (E)-(I) Tomographic reconstruction of a growing Cam2-aster processed by SIRT. See also *Video 4*. (E) Rendered view of a growing Cam2-aster from representative 120 nm thick tomography slices overlaid with a 2.4 nm tomographic slice. Centre of the Cam2-aster: yellow; microtubules: blue; tubulin sheets: green; tubulin rings: red. (F)-(I), Magnified views shown in panel (E).

**Video 3.** Cryo-tomographic reconstruction of the mature Cam2-aster. SIRT processed cryo-tomographic reconstruction of the mature Cam2-aster produced by incubation of 10 µM tubulin with 1 µM full-length CAMSAP2 for 30 min at 37 °C.

**Video 4.** Cryo-tomographic reconstruction of the growing Cam2-aster. SIRT processed cryo-tomographic reconstruction of the growing Cam2-aster produced by incubation of 10 µM tubulin with 1 µM full-length CAMSAP2 for 1 min at 37 °C.

To further investigate the Cam2-aster formation mechanism, we next applied cryo-ET to the small growing Cam2-asters formed through 1 min of incubation at 37 °C (Figure 5E-I, Video 4). We detected small condensates corresponding to the nucleation centre (growing asters) on the grid surface (yellow in Figure 5E). Short polymerizing microtubules were connected to the nucleation centres (blue in Figure 5E). Around these locations, vast numbers of small rings (red in Figure 5E and G) and sheet structures (green in Fig. 5E, H, I) were observed, although only a few rings were detectable in the mature Cam2-asters (Figure 5D). This finding suggests that rings and sheets may be important for the initial microtubule growth phase of Cam2-aster formation.

### Ring structures observed during microtubule nucleation with CAMSAP2

To understand the detailed structural mechanism of nucleation centre formation, we aimed to analyse intermediates of tubulin nucleation. We chose CC1-CKK because the centres of Cam2-asters induced by full-length CAMSAP2 were too condensed and thick for TEM to observe the inner structures. CC1-CKK exhibited similar microtubule nucleation activity with smaller condensation of the nucleation centre than full-length CAMSAP2 (Figure 4D). We analysed the time courses of microtubule nucleation and aster formation using CC1-CKK by negative stain EM (Figure 6A). Surprisingly, incubation of 10 μM tubulin with 1 μM CC1-CKK for 30 min on ice induced the formation of numerous rings with diameters of approximately 50 nm (Figure 6A), which was not observed in the tubulin-alone condition (Figure 4—figure supplement 2). After 1 min of incubation at 37 °C, many short microtubules polymerized, with slightly fewer rings than in the on-ice condition. Increasing the incubation time increased the microtubule number and length; in contrast, it further decreased the number of rings (Figure 6A and 6B). The ring number was the highest when incubation was conducted on ice, suggesting that the rings likely represent a molecular state of tubulins before their polymerization is initiated. The number started to decrease after incubation at 37 °C, and only a few rings were observed in the background after 10 min of incubation.

**Figure 6.**
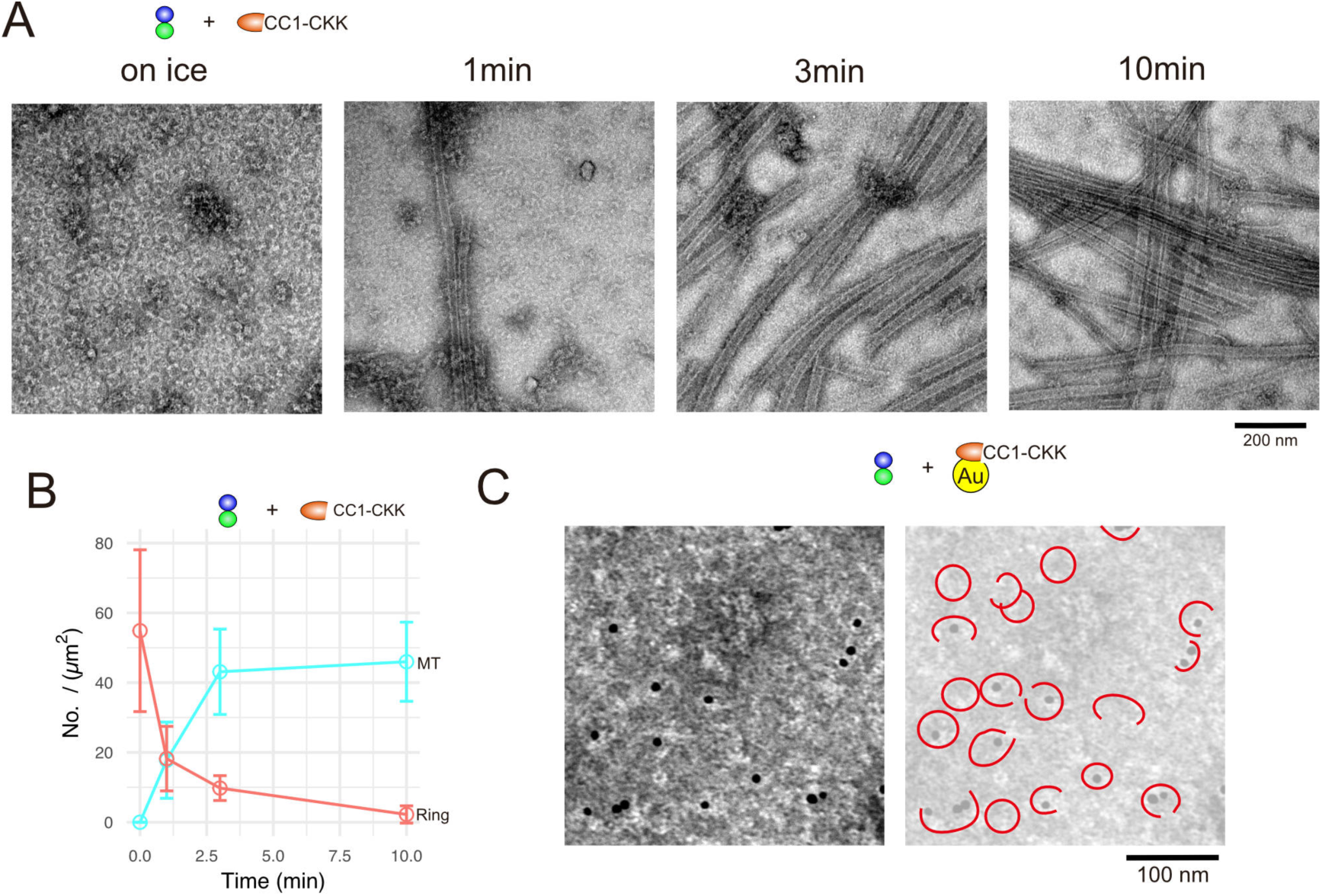
CAMSAP2 induces tubulin ring formation. (A) Negative stain EM micrographs of 10 µM tubulin polymerization with 1 µM CAMSAP2 CC1-CKK at different time points. (B) Plots of the number of tubulin rings (orange) and that of microtubules (turquoise green) at different time points (mean ± SD, from 10 independent views). (C) Negative stain EM micrographs of 10 µM tubulin with 1 µM nano-gold-labelled CAMSAP2 CC1-CKK mixed on ice for 30 min. Quality of the nano-gold labeled CAMSAP2 CC1-CKK was analyzed by negative stain EM. See Figure 6—figure supplement 1.

**Figure 6—figure supplement 1.**
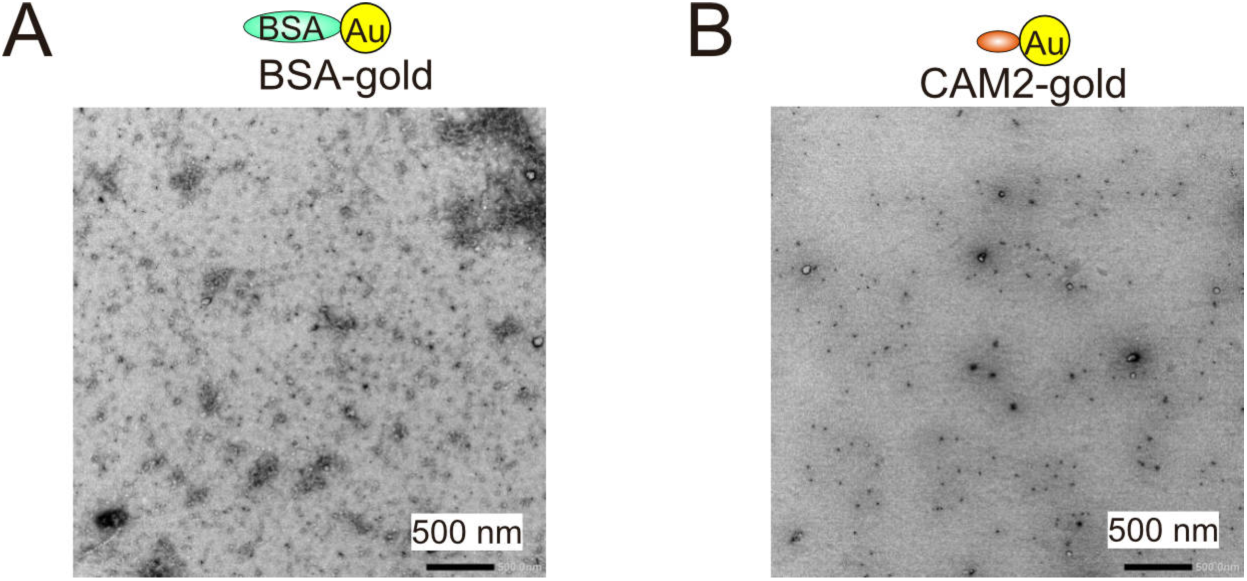
Nano-gold labelling of CAMSAP2. (A) Negative stain EM micrographs of 1 µM nano-gold-labelled BSA. (B) Negative stain EM micrographs of 1 µM nano-gold-labelled CAMSAP2 CC1-CKK.

To further examine the localization of CAMSAP2 during ring formation, we labelled CC1-CKK CAMSAP2 with nano-gold, mixed it with tubulin, and observed its localization by negative stain EM (Figure 6C). Suggestively, nano-gold densities were present at the centres of the rings or crescent-shaped tubulin oligomers (Figure 6C, black dots). It should be noted that CAMSAP2 itself was uniformly distributed and did not form any clusters on ice. Combining these results with those showing that the ring number decreased during incubation at 37 °C (Figure 6B), we can postulate that CAMSAP2 contributes to the formation of tubulin oligomers or rings and may be used as a material for initial short tubulin sheet and microtubule formation.

### CAMSAP2-induced microtubule nucleation intermediates observed by cryo-EM

To visualize the structural details of the nucleation centres during Cam2-aster growth, we used cryo-EM to observe this process at three time points. We incubated the tubulin–CC1-CKK mixture on ice for 30 min, at 37 °C for 1 min, and at 37 °C for 3 min. The concentrations of tubulin and CC1-CKK were set to 30 μM and 3 μM, respectively; the ratio was identical to that used for negative stain EM, but the concentration of each molecule was tripled to observe the rings or sheets more efficiently.

Consistent with our negative stain EM results, many tubulin rings or tubulin oligomers with crescent shapes were observed in the on-ice condition (Figure 7A and red in Figure 7B), whereas those structures were not observed in the tubulin-alone condition (Figure 7—figure supplement 1a, b). In these specimens, short tubulin sheets were also observed on ice (green in Figure 7B). Furthermore, we detected semicircular or crescent-shaped tubulin oligomers that were multiply stacked and partly flattened, suggestive of the intermediates between rings and sheets (orange in Figure 7B). Intriguingly, some densities were observed inside the rings (Figure 7C) that resembled the CAMSAP2 in the rings detected by negative stain EM with nano-gold labelling (Figure 6C). To investigate the ring structures in further detail, we performed 2D class averaging of rings from cryo-EM micrographs (Figure 7D). The rings were apparently not uniform; single rings, double rings, and spiral rings were observed. Depending on the lengths of tubulin oligomers, several forms of tubulin rings were produced. 2D classification of the rings clearly showed bumps around the ring surface corresponding to the individual tubulin monomers (Figure 7D). Weak irregular densities could still be observed in the spiral rings, which were the 2D averages of 40-100 particles. The diameters and pitches of the bumps resemble those of longitudinal protofilament rings, rather than the tubulin rings made by the lateral contacts like the γ-TuRC (Figure 7E).

**Figure 7.**
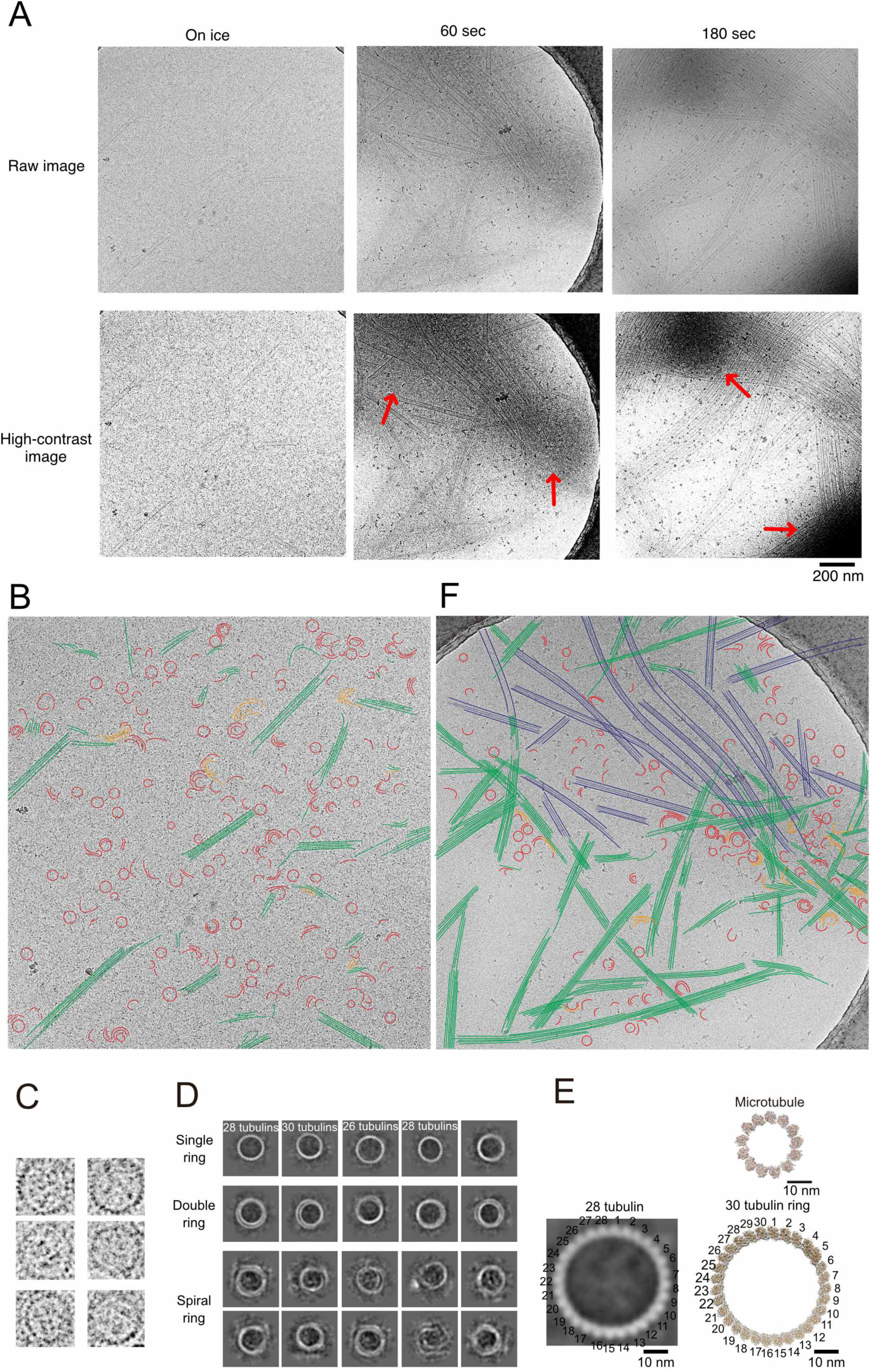
CAMSAP2 induced microtubule nucleation intermediates visualized by time-lapse cryo-EM. (A) Snapshots of growing microtubule intermediates at different time points and with different contrasts. The red arrows indicate high-density condensed areas. (B) Segmentation of the structural elements of micrographs from panel (A) (on ice). Tubulin rings: red; intermediates between ring and sheet: orange; tubulin sheets: green. (C) Ring images cropped from cryo-EM micrographs. Amorphous density could be observed in the ring. (D) 2D classification of tubulin rings. Rings of different shapes and sizes were observed, including single rings, spiral rings, and double rings. (E) Comparison of 2D average of 28 tubulin rings (left) with the single protofilament (PF) ring (bottom right) and the 13-PF microtubules seen from the plus-end (top right). The scale bars indicate 10 nm. The PF ring was generated by cropping outer microtubule ring from Microtubule-KLP10A map (EMD-7026) using its model (PDB: 6b0c) as a guide. (F) Segmentation of the structural elements of the micrographs from (A) (60 sec). Tubulin rings: red; intermediates between ring and sheet: orange; tubulin sheets: green; microtubules: blue.

**Figure 7—figure supplement 1.**
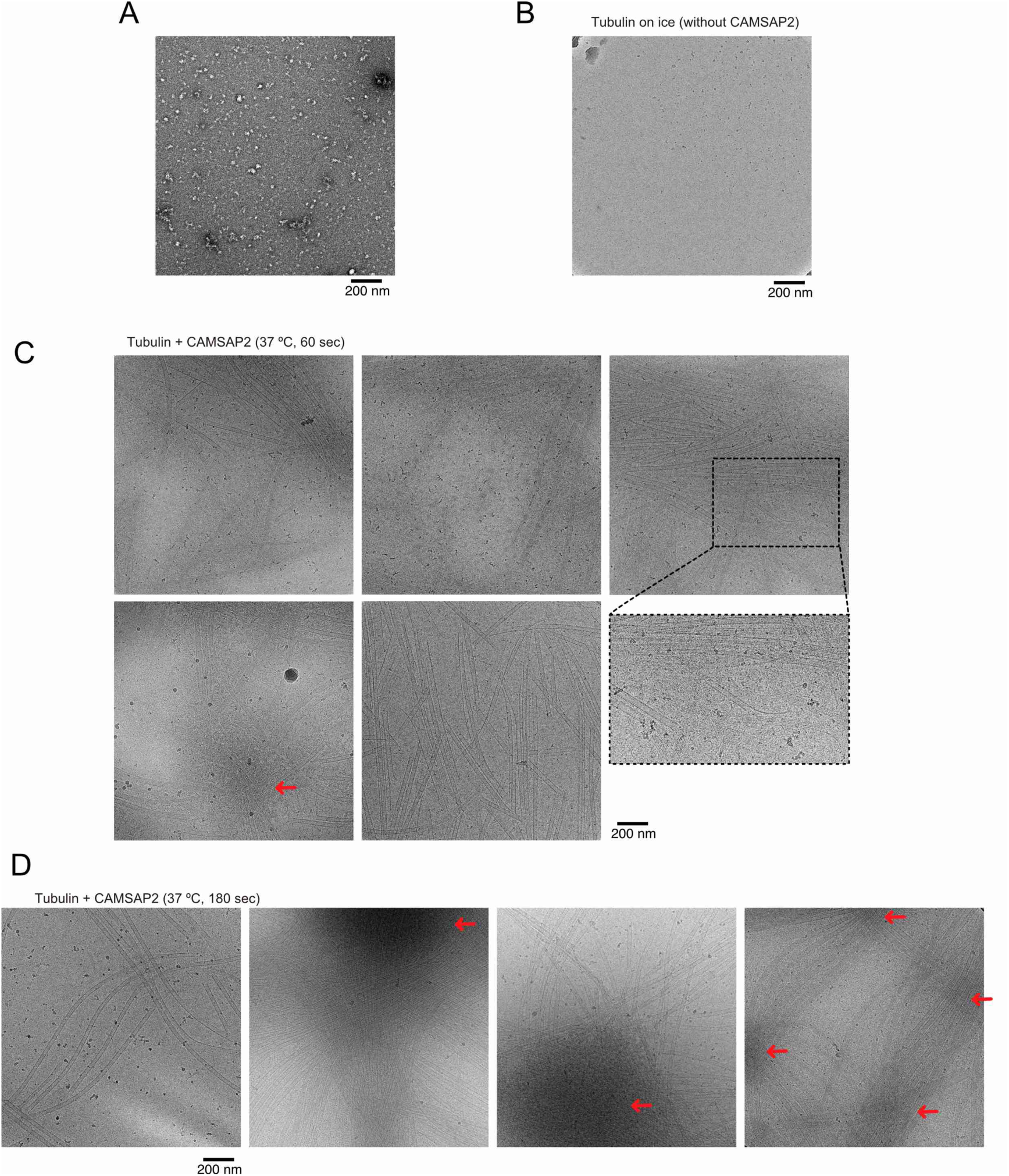
Cryo-EM visualization of Cam2-aster formation. (A) Negative stain image of 10 µM tubulin on ice. (B) Cryo-EM image of 30 µM tubulin on ice without CAMSAP2. (C) Cryo-EM images of growing microtubules with CAMSAP2 CC1-CKK after 60 sec of incubation at 37 °C. The enlarged panel shows examples of partially curved sheet-like microtubule ends, which are characteristic of growing microtubules. (D) Cryo-EM images of growing microtubules with CAMSAP2 CC1-CKK after 180 sec of incubation at 37 °C. The red arrows in panels C and D indicate high-density condensed areas.

Cryo-EM snapshots of the samples incubated at 37 °C for 1 min and 3 min clearly illustrated transient steps of microtubule nucleation and growth (Figure 7A and 7F). Before polymerization was initiated by increasing the temperature, tubulin rings, intermediates between rings and sheets, and tubulin sheets were uniformly distributed throughout the grid surface (Figure 7A). Incubation at 37 °C incubation further stimulated tubulin sheet formation; thus, the numbers and lengths of sheets were tremendously increased. After 1 minute of incubation at 37 °C, rings and short sheets tended to gather together to form a high-density condensed area (red arrow in Figure 7A, 60 sec). After 3 min of incubation, the condensates became denser, forming the central regions of Cam2-asters (red arrow in Figure 7A, 180 sec). These observations suggest that incubation at 37 °C induces the condensation of tubulin intermediates with CAMSAP2, which stimulates tubulin sheet and microtubule formation, resulting in rapid growth of Cam2-asters.

Next, we more closely investigated the structural details of the aster centres. The centres were filled with microtubule intermediates, including rings, intermediates between rings and sheets, sheets, and short microtubules (Figure 7F). Rings were also aligned along the elongating tubulin sheets that radiated from the condensates, although only a few rings were observed around mature microtubules. Some rings and sheets were assembled together, resembling the process of integration of protofilaments into tubulin sheets (orange in Figure 7F). Around the regions where numerous rings existed, the ends of microtubules or tubulin sheets often exhibited partially curved sheet-like structures that are characteristically observed for growing microtubules (Figure 7—figure supplement 1C). On the other hand, fully curved ends that are often observed for depolymerized microtubules were rarely observed.

It should be emphasized that the tubulin rings were produced on ice when no microtubules were formed. This means that the tubulin rings observed here were likely not the products of microtubule depolymerization; rather, they could be incorporated into tubulin sheets to form microtubules. We therefore conclude that CAMSAP2 facilitates protofilament ring formation and that the rings, including single rings, spiral rings, and double rings, should be nucleation intermediates that grow into microtubules through tubulin sheets that function as critical nuclei for microtubule polymerization.

## Discussion

We report here that CAMSAP2 functions as a nucleator of microtubule formation from soluble αβ-tubulin dimers. CAMSAP2 reduces the nucleation concentration barrier for microtubule polymerization to 1-2 µM, which is similar to the tubulin Cc_MT polymerization_. The acceleration of microtubule nucleation is achieved by the efficient formation of microtubule polymerization intermediates made of protofilaments inside condensed nucleation centres, which function as nucleation centres for microtubule polymerization. Each condensed region grows into the centre of a Cam2-aster, from which numerous dynamic microtubules elongate in a radial fashion. Microtubules from neighbouring asters join together to form the dense meshwork. These results suggest that CAMSAP2 contributes considerably to the fundamental step of microtubule network formation.

Our structural study indicates that CAMSAP2 induces longitudinal growth of αβ-tubulin to form a protofilament ring. Remarkably, this reaction can proceed on ice even though microtubule polymerization is not generally observed under such conditions. In the presence of CAMSAP2, tubulin ring and short sheet formation was observed on ice, and the resulting structures were distributed with uniformity. When the temperature is increased to 37 °C, tubulin rings start gathering with CAMSAP2 to form molecular clusters that will become the nucleation centres for future Cam2-asters. This molecular condensate efficiently nucleates microtubules, and is reversible and sensitive to temperature and salt concentration. In this regard, the behaviour of CAMSAP2 apparently resembles that of TPX2, the targeting protein for Xklp2 (Nashchekin et al., 2016; Wang et al., 2015). TPX2 was reported to phase separate into a co-condensate with tubulin which enhances microtubule nucleation through the spatial coordination (King and Petry, 2020). CAMSAP2 may utilize the similar strategy that should be further confirmed in the future.

In the centrosomal aster, γ-TuRC serves as a template to help αβ-tubulins align for formation of a 13-protofilament microtubule structure in a process called templated nucleation (Moritz et al., 1995; Roostalu and Surrey, 2017; Wieczorek et al., 2020; Zheng et al., 1995). Each Cam2-aster resembles the centrosomal aster, as described above, but the molecular behaviours inside the condensate, the centre of Cam2-aster, are completely different from those in the centrosomal aster. In the condensate, large numbers of tubulin rings and sheets exist, and CAMSAP2 mainly localizes inside the rings. Therefore, CAMSAP2 induces and stabilizes microtubule polymerization intermediates. Microtubules grow rapidly from the condensates, with the numbers of tubulin rings decreasing inversely (Figure 6B). Thus, the rings shown here apparently form at the rapid growing phase of microtubules, and it is reasonable to conclude that rings are used as materials to form the initial short tubulin sheets and the microtubules. However, tubulin rings have long been thought to be microtubule depolymerization products (Erickson, 1996). We could not find the differences of curvatures between the rings in the current study and previously observed depolymerized products. Indeed, McIntosh et al. also reported that electron tomography analyses showed the curvatures of protofilaments on growing and shrinking microtubules are similar (McIntosh et al., 2018). Hence, the curvatures of tubulin oligomers or the protofilaments in the growing and shrinking phases are still under debate and further high resolution structural studies on tubulin rings complexed with CAMSAP2 are necessary to confirm the functions of the rings as microtubule nucleation intermediates.

How tubulin rings are produced in the presence of CAMSAP2 remains elusive. The CKK domain of CAMSAPs has been reported to bridge two tubulin dimers in neighbouring protofilaments (Atherton et al., 2017). This binding pattern apparently does not support the single ring formation. In this respect, cryo-EM structure of TPX-2 microtubule complex provides us the possible insight (Zhang et al., 2017). TPX-2 uses two elements, ridge and wedge, to stabilize the longitudinal and the lateral contacts of microtubule. CKK of CAMSAP2 stabilizes the lateral contacts (Atherton et al., 2017). Considering that both CC3 and CKK are necessary to stimulate the microtubule nucleation, CC3 may stabilize the longitudinal contact of tubulins to support the protofilament ring formation. This hypothesis needs to be clarified by the future high resolution structural study of CAMSAP-microtubule complex.

The role of CAMSAP2 is likely to stimulate the process of spontaneous nucleation of microtubules, which is thought to start with the formation of longitudinal contacts between αβ-tubulin dimers (Roostalu and Surrey, 2017). Considering all of our data, we propose a model of spontaneous nucleation mediated by CAMSAP2 (Figure 8). First, CAMSAP2 interacts with tubulins, inducing their longitudinal stack formation until they form protofilament rings. This step can occur on ice. Incubation at 37 °C induces co-condensation of CAMSAP2 and tubulins to stimulate the efficient formation of microtubule nucleation intermediates, forming a nucleation centre. Plenty of nucleation intermediates such as rings, intermediates between ring and sheet, sheets, are produced inside the condensates. How these nucleation intermediates grow into the critical nucleus for microtubule formation is still uncertain (Erickson and Pantaloni, 1981). From the nucleation centre, many microtubules efficiently radiate outward using the nucleation intermediates as materials, producing the Cam2-aster. In this stage, CAMSAP2 also decorates microtubule lattice around the minus-end to induce efficient elongation of microtubules. Microtubules from neighbouring asters then join together and generate a microtubule meshwork among Cam2-asters, finally building the cytoskeletal framework.

**Figure 8.**
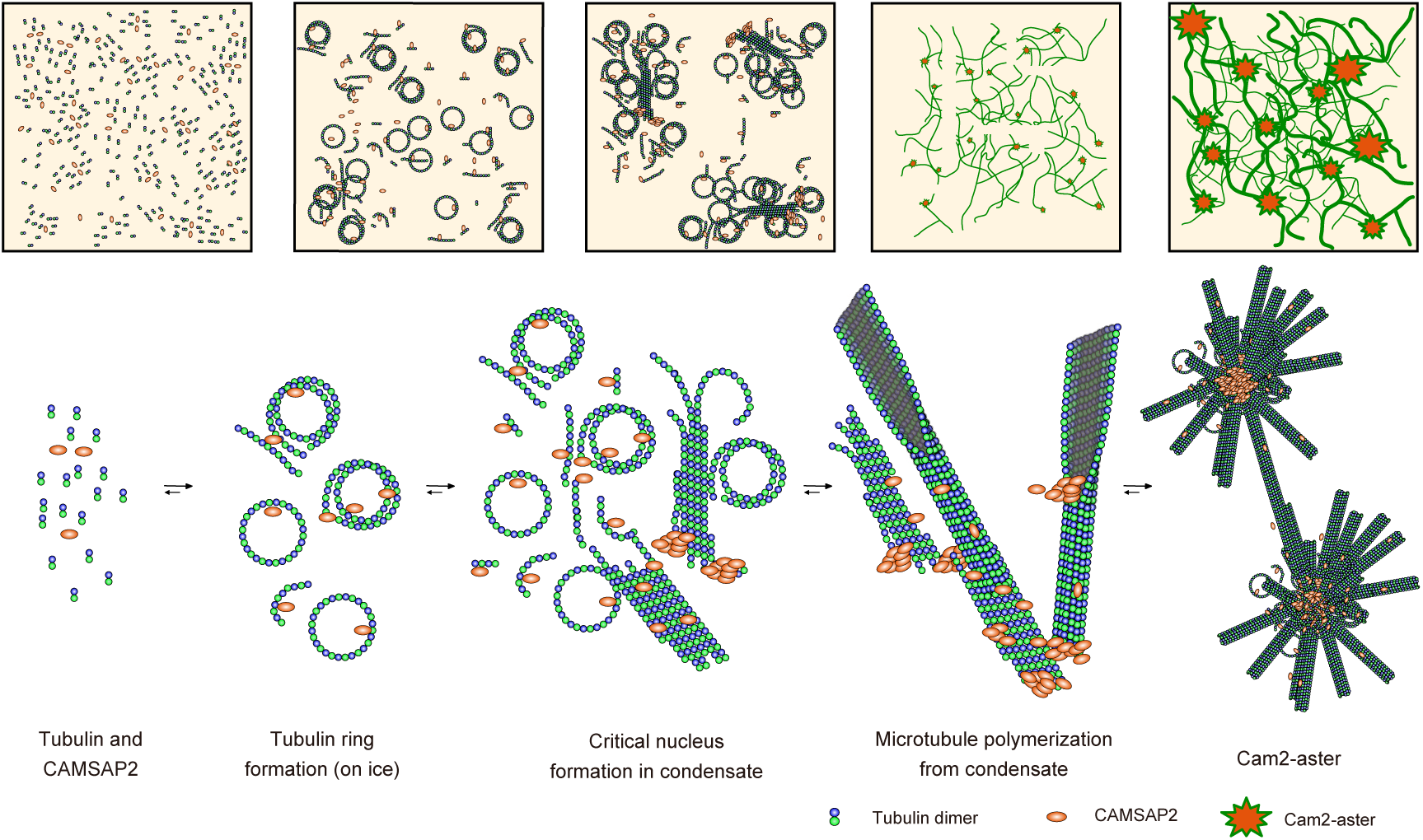
Structural model of microtubule nucleation and Cam2-aster formation induced by CAMSAP2. Structural model of tubulin nucleation, polymerization, and aster formation induced by CAMSAP2, as detailed in the main text. CAMSAP2 shifts the equilibrium to the right, as indicated by the arrow size.

Previous studies have indicated that microtubule growth occurs from CAMSAP foci (Nashchekin et al., 2016), and in differentiated epithelial cells, microtubules are anchored to the adherens junctions or apical cortices where CAMSAPs are concentrated (Toya et al., 2016). No γ-TuRC localization has been observed at these foci. Our *in vitro* data provide some insight into how CAMSAPs induce microtubule polymerization at CAMSAP-containing structures. At 50 nM, which is nearly a physiological concentration of CAMSAP proteins in cells (Liebermeister and Klipp, 2006; Wühr et al., 2014), CAMSAP2 binds to the minus-ends of microtubules but does not induce MT polymerization *in vitro* (Hendershott and Vale, 2014; Jiang et al., 2014) (Figures 1C and 3). However, we showed that in the presence of artificial CAMSAP2-containing clusters fixed on glass slides, the same concentration of CAMSAP2 could sufficiently induce microtubule polymerization from soluble tubulins *in vitro* (Figure 3). Thus, to support microtubule polymerization from particular structures, both CAMSAP-containing foci and soluble CAMSAP appear to be required.

In vertebrate cells, CAMSAPs generally decorate a single microtubule at its minus end, not inducing aster formation. Nevertheless, it has been shown that clusters of Patronin, the Drosophila homologue of CAMSAP, radiate multiple microtubules at the same time (Nashchekin et al., 2016), as observed for the Cam2-asters described here, which suggests that Cam2-aster formation may not be a purely *in vitro* event. It is possible that excess clustering of CAMSAPs and tubulins is prohibited *in vivo* through unidentified mechanisms, and CAMSAP-mediated enhancement of microtubule growth, the mechanisms of which were uncovered in this study, likely also works *in vivo* by forming a smaller or minimal unit of the CAMSAP/tubulin complex. Our findings of the interactions between CAMSAP2 and tubulins reported here using *in vitro* systems should provide insights for future studies seeking to deepen our understanding of their molecular roles in non-centrosomal microtubule network formation in living cells.

## Materials and Methods

### Tubulin preparation

Tubulin was purified from porcine brain tissue (Castoldi and Popov, 2003). The purified tubulin was labelled with 5(6)-carboxytetramethylrhodamine succinimidyl ester (TMR) (Santa Cruz Biotechnology, Dallas, TX, USA) or *N*-[6-(biotinamide)hexanoyl]-6-aminohexanoic acid N-succinimidyl ester (Biotin-LC-LC-NHS) (Tokyo Chemical Industry, Tokyo, Japan) as described previously (Al-Bassam, 2014).

### Cloning and construct design

The full-length mouse CAMSAP2 gene was PCR-amplified from the CAMSAP2-GFP plasmid (Toya et al., 2016) and subcloned between the EcoRI and XhoI restriction sites of the pFastBac1 vector (Thermofisher Scientific). GFP fusion CAMSAP2 full-length with His-Strep tag was generated from PCR-amplify genes of CAMSAP2 gene from CAMSAP2-GFP plasmid (Toya et al., 2016), GFP tag gene from CAMSAP2-GFP plasmid (Toya et al., 2016), His-Strep2-tag from 438-SNAP-V3 vector a gift from Scott Gradia (Addgene plasmid # 55223) with inserting CATCATCATCATCATCACAGCAGCGGCCTGGTGCCGCGCGGCAGCCAT sequence right in front of Strep II gene by PCR primer, and assembled into PCR amplified pFastBac vector by NEBuilder HiFi DNA assembly (NEB). CAMSAP2 mutant constructs were generated by inserting each mouse CAMSAP2 partial PCR fragment flanked by a TEV protease recognition linker (ENLYFQGSSGSSG) and a hepta-His tag at the N-terminus and C-terminus, respectively, into the pGFPS1 vector (Seki et al., 2008). This construct contains another metal-binding motif similar to the HAT tag (KDHLIHNHHKHEHAHA) at the N-terminus of the GFPS1 gene. The coding regions of all constructs were verified by Sanger sequencing.

### Protein expression

Full-length CAMSAP2 was expressed in the baculovirus expression vector system (BEVS) using insect cells. Bacmid preparation and V0 and V1 virus production were performed as described previously. The protein was expressed by Sf9 cells after infection with the V1 virus following the TEQC protocol (Imasaki et al., 2018). Other CAMSAP2 deletion constructs, summarized in Table 1, were expressed in *Escherichia. coli*. The resulting constructs were transformed into *E. coli* strain KRX (Promega) and cultured in LB medium containing 100 µg/ml ampicillin, 0.05% glucose and 0.1% rhamnose at 25 °C overnight with autoinduction. Each cell suspension was harvested into a 50 ml tube and stored at −80 °C.

**Table 1.** List of protein expression constructs used in this study

| **ID** | **construct** | **amino acids** | **vector** | **tag (N-terminal)** | **tag (C-terminal)** |
| --- | --- | --- | --- | --- | --- |
| RN82 | full length | 1–1472 | pFastBac1 | none | 6x His |
| RN63 | CC1-CKK | 696–1454 | pGFPS1 | HAT*-GFPS1-TEV* | 7x His |
| RN64 | CC3-CKK | 1157–1454 | pGFPS1 | HAT*-GFPS1-TEV* | 7x His |
| RN136 | CKK | 1332–1472 | pGFPS1 | HAT*-GFPS1-TEV* | 7x His |
| RN189 | full length | 1–1472 | pFastBac1 | His_Strep2-3Csite | none |

\*These tags carry modifications. See the Methods for details.

### Protein purification

For purification of the *E. coli*-expressed constructs, we applied the protocol described below. *E. coli* cells were thawed, resuspended in lysis buffer (40 mM Na-phosphate buffer (pH 7.4), 400 mM NaCl, 5 mM imidazole, 14 mM β-mercaptoethanol, 0.7 µM leupeptin, 2 µM pepstatin A, 1 mM PMSF, and 2 mM benzamidine), and lysed by sonication. The lysate was clarified by centrifugation (16,000 ×*g*), and the supernatant was applied to a HIS-Select Nickel Affinity Gel column (Sigma) equilibrated in lysis buffer. The column was washed with ∼10 column volumes of lysis buffer containing 5 mM imidazole. His-tagged proteins were eluted with ∼4 column volumes of lysis buffer containing 150 mM imidazole. The elution fractions were pooled and diluted with water to ∼150 mM NaCl for cation exchange chromatography. The diluted solutions were clarified by centrifugation using an Avanti JXN30 centrifuge (Beckman Coulter) with a JA30.50 Ti rotor at 20,000 ×*g* and applied to a HiTrap SP HP column (GE Healthcare; 1 ml) equilibrated in buffer A (20 mM Na-phosphate buffer (pH 7.0), 1 mM Na-EDTA, and 1 mM DTT) containing 150 mM NaCl. The column was washed with the same buffer until a stable baseline was reached. Proteins were eluted with a 20 ml linear gradient from 150 mM to 500 mM at a flow rate of 0.5 ml/min using an ÄKTA Pure instrument (GE Healthcare). The fractions containing the target proteins were pooled and concentrated to less than 0.5 ml using an Amicon Ultra concentrator (Merck Millipore; 10 kDa MWCO). The concentrate was applied to a Superdex 200 Increase 10/300 GL size exclusion chromatography column (GE Healthcare) equilibrated in sizing buffer (20 mM HEPES-KOH pH 7.5, 300 mM NaCl, and 1 mM DTT) at a flow rate of 0.4 ml/min. The peak fractions were pooled, concentrated with an Amicon Ultra 10 kDa MWCO concentrator, flash-frozen in liquid nitrogen and stored at −80 °C until use.

For constructs expressed in insect cells through BEVS, cells were thawed, resuspended in BEVS lysis buffer containing 50 mM HEPES-KOH pH 7.5, 400 mM KCl, 5 mM imidazole, 10% glycerol, 5 mM β-mercaptoethanol, 0.7 µM leupeptin, 2 µM pepstatin A, 1 mM PMSF, and 2 mM benzamidine, and lysed by sonication. The lysate was centrifuged using an Avanti JXN30 (Beckman Coulter) with a JA30.50 Ti rotor at 28,000 rpm (94,830 ×*g*), and the supernatant was applied to a HIS-Select Nickel Affinity Gel column (Sigma) equilibrated in lysis buffer. The column was washed by passing through 20 resin volumes of lysis buffer. The bound proteins were eluted with lysis buffer containing 300 mM imidazole. The eluate was concentrated by an Amicon Ultra 10 kDa MWCO concentrator and applied to a Superose6 Increase 10/300 GL size exclusion chromatography column equilibrated in sizing buffer (50 mM HEPES-KOH pH 7.5, 400 mM KCl, 10% glycerol, and 5 mM DTT). The peak fractions were pooled, concentrated by an Amicon Ultra 10 kDa MWCO concentrator, flash-frozen in liquid nitrogen, and stored at −80 °C. The quality of the samples was assessed by SDS-PAGE (Figure 1B, C).

### Protein conjugation of gold nanoparticles

Twenty-three microlitres of a 10-nm diameter gold nanoparticle suspension (Sigma cat. No. 752584) was mixed with 1 µl of 100 mM Tris-HCl pH 8.5 to adjust the pH for conjugation. The purified protein or BSA in parallel control experiments was diluted to 15 µM in sizing buffer containing 1 M KCl. Six microlitres of the diluted protein solution was added slowly to the gold nanoparticle suspension under constant mixing by gentle vortexing. The mixture was left on ice for 30 min and subjected to the negative stain EM analysis. No noticeable colour change of the conjugate suspension, which would indicate aggregation, was observed during the experiments.

### Spontaneous nucleation assay (pelleting assay)

The method was based on that in Wieczorek et al. with some modifications (Wieczorek et al., 2015). For the assay testing tubulin alone, the indicated concentrations of tubulin in PEM buffer supplemented with 1 mM GTP (PEM GTP) were incubated on ice for 30 min and then at 37 °C for 30 min. Total samples for SDS-PAGE were prepared at this stage. These samples were centrifuged in a TOMY MX-307 for 30 min at 15,000 rpm and 35 °C. The supernatant was discarded, and PEM GTP was supplemented for washing, centrifugation, and recovery of the pellet. The pellets were resuspended in cold PEM GTP, incubated on ice for 30 min for depolymerization, and centrifuged for 30 min at 15,000 rpm and 4 °C to remove aggregates or debris. The supernatants were resolved on SDS-PAGE gels, and the bands were quantified by FIJI (ImageJ) (Schneider et al., 2012) using a standard made from tubulin aliquots loaded at 1, 2, 3, and 5 µM. For tubulin with CAMSAP2, an assay was performed following the methods of the tubulin-only pelleting assay except that 1 µM CAMSAP2 was added at the initial polymerization step, and CAMSAP2 sizing buffer was used for depolymerization.

### Preparation of GMPCPP seeds

TAMRA-tubulin (2.5 μM) and biotin-LC-LC-tubulin (2.5 μM) were polymerized in 100 μl of PEM buffer supplemented with 1 mM GMPCPP and 1 mM DTT at 37 °C for 20 min. Equal amounts of prewarmed PEM buffer were added, and the mixtures were ultracentrifuged at 80,000 rpm for 5 min at 30 °C using a TLA100 rotor and an Optima TLX ultracentrifuge (Beckman). The microtubule pellets were resuspended in PEM supplemented with 1 mM DTT. The GMPCPP seeds were aliquoted into PCR tubes (NIPPON Genetics, Tokyo, Japan), snap-frozen in liquid N_2_ and stored at −80 °C. The GMPCPP seeds were quickly thawed at 37 °C immediately before use.

### Binding of CAMSAP2 on dynamic microtubules

Coverslips (#1.5H thickness, 22 x 22 mm; Thorlabs Japan, Tokyo, Japan) were washed by ultrasonic bath in 1 M HCl (Fujifilm Wako, Tokyo, Japan) for 3 hours, in deionized distilled water (DDW) for 3 hours and 1 hour, in 70% ethanol for 1 hour and then in 99.5% ethanol for 1 hour. Flow chambers were prepared at room temperature. The flow chambers were constructed with double-sided tape (Nichiban, Tokyo, Japan). The volume of each flow chamber was approximately 15 μl. The chambers were incubated with 0.5 mg/ml PLL-PEG-biotin (SuSoS, Dübendorf, Switzerland) for 10 min followed by 0.5 mg/ml streptavidin (Fujifilm Wako, Tokyo, Japan) for 2 min. The GMPCPP seeds were incubated for 5 min and washed with 50 μl of assay buffer (100 mM PIPES pH 6.8, 100 mM KCl, 2 mM MgCl_2_, 1 mM EGTA, 0.5% Pluronic F-127, 0.1 mg/ml Biotin-BSA, 0.2 mg/ml κ-casein). Then, 10 μM tubulin, 0.5 μM TAMRA-tubulin and 50 nM CAMSAP2 in assay buffer supplemented with 10 nM protocatechuate-3,4-dioxygenase (PCD) (Oriental Yeast, Tokyo, Japan), 2.5 mM protocatechuic acid (PCA) (Fujifilm Wako) and 1 mM Trolox (Fujifilm Wako) was added into the chamber and observed by total internal reflection fluorescent (TIRF) microscopy at 37 °C. The TIRF system consisted of a Ti2E microscope (Nikon, Tokyo, Japan) equipped with a Ti2-LAPP TIRF system (Nikon), an iXon Life 897 EMCCD camera (Andor), a CFI Apochromat TIRF lens (NA 1.49, x100) (Nikon) and a heating stage (Tokai Hit, Fujinomiya, Japan). Images were obtained every 5 seconds for 20 min.

### Time-lapse imaging of CAMSAP2-dependent aster formation

PLL-PEG-biotin coating was performed as described above. Instead of GMPCPP MTs, a biotinylated anti-GFP antibody (MBL, Tokyo, Japan) diluted 1:10 in PEM buffer supplemented with 100 mM KCl was added into the chamber and incubated for 2 min. The chamber was washed with 50 μl of assay buffer. CAMSAP2-GFP (0.5 μM) diluted in assay buffer was incubated for 5 min, and the chamber was washed with 50 μl of assay buffer again. Next, 10 μM tubulin, 0.5 μM TAMRA-tubulin and the indicated concentrations of CAMSAP2 in assay buffer supplemented with 10 nM PCD, 2.5 mM PCA and 1 mM Trolox were added into the chamber and were observed by TIRF microscopy as described above. Observation was performed at 37 °C. Images were obtained every 5 seconds for 20 min.

### Grid preparation and data collection for negative stain electron microscopy

Samples, including tubulin, CAMSAP2, and/or CAMSAP2 mutant samples, were diluted with PEM buffer (100 mM PIPES pH 6.8, 1 mM MgCl_2_, 1 mM EGTA, and 1 mM GTP) and incubated on ice for 30 min before blotting. The copolymerized samples were mixed, incubated on ice for 30 min, and then transferred to 37 °C for 1, 3, 10, or 30 min. After incubation, 4 µl of each sample was applied to a glow-discharged carbon-coated 200-mesh Cu grid (EM Japan). Any excess solution was wicked off with filter paper as soon as possible. The grid was stained with 2% uranyl acetate, the excess stain was blotted with filter paper, and the grid was air-dried. The specimens were observed using a JEM-1400Plus electron microscope (JEOL) operated at 120 kV and equipped with a JEOL Matataki CMOS camera at nominal magnifications of 600-, 10,000-, 20,000-, and 40,000-fold.

### Grid preparation for cryo-electron microscopy

Next, 30 μM tubulin and 3 μM full-length CAMSAP2 or CC1-CKK in PEM buffer (100 mM PIPES-KOH pH 6.8, 1 mM MgCl_2_, 1 mM EGTA, and 1 mM GTP) were mixed and incubated on ice for 30 min. For induction of microtubule polymerization, the samples were further incubated at 37 °C for 1 min or at 37 °C for 3 min. All samples were placed onto glow-discharged R2/2 Quantifoil 300-mesh Cu grids with carbon support film, blotted with filter paper (Whatman No. 1) for 2 sec, and plunge-frozen in liquid ethane with a Vitrobot Mark IV (Thermofisher Scientific).

### Cryo-electron microscopy

Data acquisition was performed by using a Glacios Cryo Transmission Electron Microscope operated at 200 kV with a Falcon III EC Direct Electron Detector (Thermofisher Scientific) at 28,000-fold nominal magnification. A total of 176 micrographs were collected with a total dose of 20 e^-^/Å^2^ at a pixel size of 3.57 Å. For the 2D classification, 5,881 particles in total were manually picked from 89 micrographs and subjected to two rounds of reference-free 2D classification in cryoSPARC (Punjani et al., 2017).

### Cryo-electron tomography

A Titan Krios 300 kV transmission electron microscope equipped with a Falcon II Direct Electron Detector (Thermofisher Scientific) at nominal magnifications of 11,000-fold (for mature asters) and 14,000-fold (for growing asters) was used for 3D structure analysis. A cryo-ET tilt series was recorded with TEM Tomography Data Acquisition software. The dose for one image was calculated as 1.5 electrons/Å^2^. The tilt series was obtained using a Saxton scheme with steps of 3 degrees up to ±70 degrees. Sixty-seven micrographs were taken for one series; the total dose was approximately 100 e^-^/Å^2^ at pixel sizes of 7.4 Å (for mature asters) and 5.9 Å (for growing asters). The tilt series images were reconstructed into 3D tomograms by using the IMOD software package (Kremer et al., 1996). The micrographs were aligned by cross-correlation and then aligned by tracking 20 nm gold fiducial beads coated on the grid. The aligned micrographs were reconstructed for visual analysis using IMOD SIRT. CTF correction was not performed. Particles from exemplary classes are displayed in Figure 6E-I.

## Acknowledgements

We thank R.J. McKenney at UC Davis for providing the full-length GFP-CAMSAP2 construct and N. Kajimura for helping with the electron microscopy data collection at the Research Center for Ultra-High-Voltage Electron Microscopy (Nanotechnology Open Facilities) at Osaka University. We are also grateful to K. Ikegami for his critical reading of the manuscript. We thank K. Chin for her research management support. This work was supported by the Nanotechnology Platform of the MEXT, Japan; the RIKEN Pioneering Project “Dynamic Structural Biology”; and the Platform Project for Supporting Drug Discovery and Life Science Research (Basis for Supporting Innovative Drug Discovery and Life Science Research (BINDS)) from AMED under Grant Number JP18am0101082. We acknowledge support from the Japan Society for the Promotion of Science (KAKENHI; 19K07246 to T. I., 25221104 to M. T., and 15K08168 to R. N.), AMED-CREST from the Japan Agency for Medical Research and Development (1005341 to R. N.), the Japan Science and Technology Agency/PRESTO (JPMJPR14L2 to T. I.), the Takeda Science Foundation to T. I. and R. N, the Mochida Memorial Foundation for Medical and Pharmaceutical Research to T. I. and R. N., the Uehara Memorial Foundation to R. N., Bristol-Myers Squibb to R. N., and the Hyogo Science and Technology Association to R. N.

## Contributions

T. I., S. N., M. S., M. T. and R. N. conceived the project. T. I., S. K., Y. S. H., M. A., A. S., Y. T., N. S., S. T., Y. Y., T. S., Y. S., T. S. and E. N. performed DNA construction, biochemical analyses, and cell experiments. S. N. performed TIRF assays. T. I., H. S., K. A., K. M. and R. N. performed the cryo-EM data collection and analysis. T. I., S. K., S. N., and R. N. prepared the figures. All authors discussed the results, and T. I., S. K., S. N., M. T. and R. N. wrote the manuscript.

## Corresponding author

Correspondence and requests for materials should be addressed to.

## References

Akhmanova A, Steinmetz MO. 2019. Microtubule minus-end regulation at a glance. J Cell Sci 132:jcs227850. doi:10.1242/jcs.227850

Akhmanova A, Steinmetz MO. 2015. Control of microtubule organization and dynamics: two ends in the limelight. Nat Rev Mol Cell Biol 16:711–726. doi:10.1038/nrm4084

Akhmanova A, Steinmetz MO. 2008. Tracking the ends: a dynamic protein network controls the fate of microtubule tips. Nat Rev Mol Cell Biol 9:309–322. doi:10.1038/nrm2369

Al-Bassam J. 2014. Reconstituting Dynamic Microtubule Polymerization Regulation by TOG Domain ProteinsMethods in Enzymology. Elsevier. pp. 131–148. doi:10.1016/B978-0-12-397924-7.00008-X

Alushin GM, Lander GC, Kellogg EH, Zhang R, Baker D, Nogales E. 2014. High-Resolution Microtubule Structures Reveal the Structural Transitions in αβ-Tubulin upon GTP Hydrolysis. Cell 157:1117–1129. doi:10.1016/j.cell.2014.03.053

Atherton J, Jiang K, Stangier MM, Luo Y, Hua S, Houben K, van Hooff JJE, Joseph A-P, Scarabelli G, Grant BJ, Roberts AJ, Topf M, Steinmetz MO, Baldus M, Moores CA, Akhmanova A. 2017. A structural model for microtubule minus-end recognition and protection by CAMSAP proteins. Nat Struct Mol Biol 24:931–943. doi:10.1038/nsmb.3483

Atherton J, Luo Y, Xiang S, Yang C, Rai A, Jiang K, Stangier M, Vemu A, Cook AD, Wang S, Roll-Mecak A, Steinmetz MO, Akhmanova A, Baldus M, Moores CA. 2019. Structural determinants of microtubule minus end preference in CAMSAP CKK domains. Nat Commun 10:5236. doi:10.1038/s41467-019-13247-6

Castoldi M, Popov AV. 2003. Purification of brain tubulin through two cycles of polymerization–depolymerization in a high-molarity buffer. Protein Expr Purif 32:83–88. doi:10.1016/S1046-5928(03)00218-3

Chuang M, Goncharov A, Wang S, Oegema K, Jin Y, Chisholm AD. 2014. The Microtubule Minus-End-Binding Protein Patronin/PTRN-1 Is Required for Axon Regeneration in C. elegans. Cell Rep 9:874–883. doi:10.1016/j.celrep.2014.09.054

Dammermann A, Desai A, Oegema K. 2003. The minus end in sight. Curr Biol 13:R614–R624. doi:10.1016/S0960-9822(03)00530-X

Desai A, Mitchison TJ. 1997. MICROTUBULE POLYMERIZATION DYNAMICS. Annu Rev Cell Dev Biol 13:83–117. doi:10.1146/annurev.cellbio.13.1.83

Erickson HP. 1996. Protofilaments and rings, two conformations of the tubulin family conserved from bacterial FtsZ to alpha/beta and gamma tubulin. J Cell Biol 135:5–8. doi:10.1083/jcb.135.1.5

Erickson HP, Pantaloni D. 1981. The role of subunit entropy in cooperative assembly. Nucleation of microtubules and other two-dimensional polymers. Biophys J 34:293–309. doi:10.1016/S0006-3495(81)84850-3

Goodwin SS, Vale RD. 2010. Patronin Regulates the Microtubule Network by Protecting Microtubule Minus Ends. Cell 143:263–274. doi:10.1016/j.cell.2010.09.022

Hannak E, Oegema K, Kirkham M, Gönczy P, Habermann B, Hyman AA. 2002. The kinetically dominant assembly pathway for centrosomal asters in Caenorhabditis elegans is γ-tubulin dependent. J Cell Biol 157:591–602. doi:10.1083/jcb.200202047

Hendershott MC, Vale RD. 2014. Regulation of microtubule minus-end dynamics by CAMSAPs and Patronin. Proc Natl Acad Sci 111:5860–5865. doi:10.1073/pnas.1404133111

Howard J, Hyman AA. 2003. Dynamics and mechanics of the microtubule plus end. Nature 422:753–758. doi:10.1038/nature01600

Imasaki T, Wenzel S, Yamada K, Bryant ML, Takagi Y. 2018. Titer estimation for quality control (TEQC) method: A practical approach for optimal production of protein complexes using the baculovirus expression vector system. PLOS ONE 13:e0195356. doi:10.1371/journal.pone.0195356

Jiang K, Faltova L, Hua S, Capitani G, Prota AE, Landgraf C, Volkmer R, Kammerer RA, Steinmetz MO, Akhmanova A. 2018. Structural Basis of Formation of the Microtubule Minus-End-Regulating CAMSAP-Katanin Complex. Structure 26:375–382.e4. doi:10.1016/j.str.2017.12.017

Jiang K, Hua S, Mohan R, Grigoriev I, Yau KW, Liu Q, Katrukha EA, Altelaar AFM, Heck AJR, Hoogenraad CC, Akhmanova A. 2014. Microtubule Minus-End Stabilization by Polymerization-Driven CAMSAP Deposition. Dev Cell 28:295–309. doi:10.1016/j.devcel.2014.01.001

King MR, Petry S. 2020. Phase separation of TPX2 enhances and spatially coordinates microtubule nucleation. Nat Commun 11:270. doi:10.1038/s41467-019-14087-0

Kremer JR, Mastronarde DN, McIntosh JR. 1996. Computer Visualization of Three-Dimensional Image Data Using IMOD. J Struct Biol 116:71–76. doi:10.1006/jsbi.1996.0013

Kuchnir Fygenson D, Flyvbjerg H, Sneppen K, Libchaber A, Leibler S. 1995. Spontaneous nucleation of microtubules. Phys Rev E 51:5058–5063. doi:10.1103/PhysRevE.51.5058

Liebermeister W, Klipp E. 2006. Bringing metabolic networks to life: integration of kinetic, metabolic, and proteomic data. Theor Biol Med Model 3:42. doi:10.1186/1742-4682-3-42

Marcette JD, Chen JJ, Nonet ML. 2014. The Caenorhabditis elegans microtubule minus-end binding homolog PTRN-1 stabilizes synapses and neurites. eLife 3:e01637. doi:10.7554/eLife.01637

Martin M, Veloso A, Wu J, Katrukha EA, Akhmanova A. 2018. Control of endothelial cell polarity and sprouting angiogenesis by non-centrosomal microtubules. eLife 7:e33864. doi:10.7554/eLife.33864

McIntosh JR, O’Toole E, Morgan G, Austin J, Ulyanov E, Ataullakhanov F, Gudimchuk N. 2018. Microtubules grow by the addition of bent guanosine triphosphate tubulin to the tips of curved protofilaments. J Cell Biol 217:2691–2708. doi:10.1083/jcb.201802138

Meng W, Mushika Y, Ichii T, Takeichi M. 2008. Anchorage of Microtubule Minus Ends to Adherens Junctions Regulates Epithelial Cell-Cell Contacts. Cell 135:948–959. doi:10.1016/j.cell.2008.09.040

Moritz M, Braunfeld MB, Sedat JW, Albertst B, Agard DA. 1995. Microtubule nucleation by γ-tubulin-containing rings in the centrosome. Nature 378:638– 640.

Nashchekin D, Fernandes AR, St Johnston D. 2016. Patronin/Shot Cortical Foci Assemble the Noncentrosomal Microtubule Array that Specifies the Drosophila Anterior-Posterior Axis. Dev Cell 38:61–72. doi:10.1016/j.devcel.2016.06.010

Nogales E, Whittaker M, Milligan RA, Downing KH. 1999. High-Resolution Model of the Microtubule. Cell 96:79–88. doi:10.1016/S0092-8674(00)80961-7

Noordstra I, Liu Q, Nijenhuis W, Hua S, Jiang K, Baars M, Remmelzwaal S, Martin M, Kapitein LC, Akhmanova A. 2016. Control of apico–basal epithelial polarity by the microtubule minus-end-binding protein CAMSAP3 and spectraplakin ACF7. J Cell Sci 129:4278–4288. doi:10.1242/jcs.194878

O’Toole E, Greenan G, Lange KI, Srayko M. 2012. The Role of γ-Tubulin in Centrosomal Microtubule Organization. PLoS ONE 7:e29795. doi:10.1371/journal.pone.0029795

Pongrakhananon V, Saito H, Hiver S, Abe T, Shioi G, Meng W, Takeichi M. 2018. CAMSAP3 maintains neuronal polarity through regulation of microtubule stability. Proc Natl Acad Sci 115:9750–9755. doi:10.1073/pnas.1803875115

Punjani A, Rubinstein JL, Fleet DJ, Brubaker MA. 2017. cryoSPARC: algorithms for rapid unsupervised cryo-EM structure determination. Nat Methods 14:290– 296. doi:10.1038/nmeth.4169

Richardson CE, Spilker KA, Cueva JG, Perrino J, Goodman MB, Shen K. 2014. PTRN-1, a microtubule minus end-binding CAMSAP homolog, promotes microtubule function in Caenorhabditis elegans neurons. eLife 3:e01498. doi:10.7554/eLife.01498

Roostalu J, Cade NI, Surrey T. 2015. Complementary activities of TPX2 and chTOG constitute an efficient importin-regulated microtubule nucleation module. Nat Cell Biol 17:1422–1434. doi:10.1038/ncb3241

Roostalu J, Surrey T. 2017. Microtubule nucleation: beyond the template. Nat Rev Mol Cell Biol 18:702–710. doi:10.1038/nrm.2017.75

Sampaio P, Rebollo E, Varmark H, Sunkel CE. 2001. Organized microtubule arrays in γ-tubulin-depleted Drosophila spermatocytes. Curr Biol 11:1788–1793.

Schneider CA, Rasband WS, Eliceiri KW. 2012. NIH Image to ImageJ: 25 years of image analysis. Nat Methods 9:671–675. doi:10.1038/nmeth.2089

Seki E, Matsuda N, Yokoyama S, Kigawa T. 2008. Cell-free protein synthesis system from Escherichia coli cells cultured at decreased temperatures improves productivity by decreasing DNA template degradation. Anal Biochem 377:156–161. doi:10.1016/j.ab.2008.03.001

Tanaka N, Meng W, Nagae S, Takeichi M. 2012. Nezha/CAMSAP3 and CAMSAP2 cooperate in epithelial-specific organization of noncentrosomal microtubules. Proc Natl Acad Sci 109:20029–20034. doi:10.1073/pnas.1218017109

Toya M, Kobayashi S, Kawasaki M, Shioi G, Kaneko M, Ishiuchi T, Misaki K, Meng W, Takeichi M. 2016. CAMSAP3 orients the apical-to-basal polarity of microtubule arrays in epithelial cells. Proc Natl Acad Sci 113:332–337. doi:10.1073/pnas.1520638113

Voter WA, Erickson HP. 1984. The Kinetics of Microtubule Assembly. J Biol Chem 259:10430–10438.

Wang S, Wu D, Quintin S, Green RA, Cheerambathur DK, Ochoa SD, Desai A, Oegema K. 2015. NOCA-1 functions with γ-tubulin and in parallel to Patronin to assemble non-centrosomal microtubule arrays in C. elegans. eLife 4:e08649. doi:10.7554/eLife.08649

Wieczorek M, Bechstedt S, Chaaban S, Brouhard GJ. 2015. Microtubule-associated proteins control the kinetics of microtubule nucleation. Nat Cell Biol 17:907– 916. doi:10.1038/ncb3188

Wieczorek M, Urnavicius L, Ti S-C, Molloy KR, Chait BT, Kapoor TM. 2020. Asymmetric Molecular Architecture of the Human γ-Tubulin Ring Complex. Cell 180:165–175. doi:10.1016/j.cell.2019.12.007

Wu J, de Heus C, Liu Q, Bouchet BP, Noordstra I, Jiang K, Hua S, Martin M, Yang C, Grigoriev I, Katrukha EA, Altelaar AFM, Hoogenraad CC, Qi RZ, Klumperman J, Akhmanova A. 2016. Molecular Pathway of Microtubule Organization at the Golgi Apparatus. Dev Cell 39:44–60. doi:10.1016/j.devcel.2016.08.009

Wühr M, Freeman RM, Presler M, Horb ME, Peshkin L, Gygi SP, Kirschner MW. 2014. Deep Proteomics of the Xenopus laevis Egg using an mRNA-Derived Reference Database. Curr Biol 24:1467–1475. doi:10.1016/j.cub.2014.05.044

Yau KW, van Beuningen SFB, Cunha-Ferreira I, Cloin BMC, van Battum EY, Will L, Schätzle P, Tas RP, van Krugten J, Katrukha EA, Jiang K, Wulf PS, Mikhaylova M, Harterink M, Pasterkamp RJ, Akhmanova A, Kapitein LC, Hoogenraad CC. 2014. Microtubule Minus-End Binding Protein CAMSAP2 Controls Axon Specification and Dendrite Development. Neuron 82:1058– 1073. doi:10.1016/j.neuron.2014.04.019

Zhang R, Roostalu J, Surrey T, Nogales E. 2017. Structural insight into TPX2-stimulated microtubule assembly. eLife 6:e30959. doi:10.7554/eLife.30959

Zheng Y, Wong ML, Alberts B. 1995. Nucleation of microtubule assembly by a γ-tubulin-containing ring complex. Nature 378:578–583.

